# An astrocytic signaling loop for frequency-dependent control of dendritic integration and spatial learning

**DOI:** 10.1101/2021.11.05.467400

**Authors:** Kirsten Bohmbach, Nicola Masala, Eva M. Schönhense, Katharina Hill, André N. Haubrich, Andreas Zimmer, Thoralf Opitz, Heinz Beck, Christian Henneberger

## Abstract

Dendrites of hippocampal CA1 pyramidal cells amplify clustered glutamatergic input by activation of voltage-gated sodium channels and N-methyl-D-aspartate receptors (NMDARs). NMDAR activity depends on the presence of NMDAR co-agonists such as D-serine, but how co-agonists influence dendritic integration is not well understood. Using combinations of whole-cell patch clamp, iontophoretic glutamate application, two-photon excitation fluorescence microscopy and glutamate uncaging we found that exogenous D-serine reduces the threshold of dendritic spikes and increases their amplitude. Triggering an astrocytic mechanism controlling endogenous D-serine supply via endocannabinoid receptors (CBRs) also increased dendritic spiking. Unexpectedly, this pathway was activated by pyramidal cell activity primarily in the theta range, which required HCN channels and astrocytic CB1Rs. Therefore, astrocytes close a positive and frequency-dependent feedback loop between pyramidal cell activity and their integration of dendritic input. Its disruption led to an impairment of spatial memory, which demonstrates its behavioral relevance.

## Introduction

Dendrites are the main neuronal input structure, receiving thousands of excitatory and inhibitory inputs. Their integration of incoming synaptic input is determined by passive and active mechanisms (for review Stuart and Spruston 2015). In addition to affecting voltage propagation, voltage-gated mechanisms can strongly amplify synchronous and spatially clustered synaptic glutamatergic inputs through generation of dendritic spikes. In dendrites of CA1 pyramidal cells of the hippocampus, this involves for instance voltage-dependent sodium channels and glutamate receptors of the N-methyl-D-aspartate subtype (NMDARs) (Ariav et al. 2003; Grienberger et al. 2014; Harnett et al. 2012; Losonczy and Magee 2006). Computationally, this represents, for example, a mechanism of coincidence detection (Stuart and Spruston 2015) and numerous studies have described functional roles for dendritic spikes in CA1 pyramidal cells. For instance, dendritic spike generation plays a role for synaptic long-term plasticity in vitro (Remy and Spruston 2007) and dendritic spikes are strongly associated with somatic complex spike bursts in vivo (Grienberger et al. 2014). The latter have been shown to occur in hippocampal place cells and to be spatially tuned *in vivo* (Epsztein et al. 2011; Harvey et al. 2009). More recently, studies using two-photon imaging of dendritic Ca^2+^ signals in behaving mice have directly visualized how dendritic Ca^2+^ spikes can participate in the representation and formation of place fields (Sheffield et al. 2017; Sheffield and Dombeck 2015).

The activation of NMDARs during dendritic spiking is driven by the binding of their ligand glutamate and postsynaptic depolarization. However, opening of NMDARs also requires the binding of a co-agonist, either glycine or D-serine (Johnson and Ascher 1987; Kleckner and Dingledine 1988). At glutamatergic synapses of CA1 pyramidal cells, the level of saturation of the NMDAR co-agonist binding site with co-agonists can vary in an activity-dependent manner (Adamsky et al. 2018; Henneberger et al. 2010; Papouin et al. 2017; Robin et al. 2018; Tsai et al. 2004). This indicated to us that the generation and properties of dendritic spikes should be controlled by the availability of NMDAR co-agonists and its dynamic changes.

Between the two NMDAR co-agonists, the regulation of extracellular D-serine levels has recently attracted considerable attention (Adamsky et al. 2018; Henneberger et al. 2010; Le Bail et al. 2015; Papouin et al. 2017; Robin et al. 2018). We therefore asked if D-serine and documented mechanisms that control D-serine levels affect dendritic integration of CA1 pyramidal cell and spatial memory formation. We found that this was indeed the case and revealed an unexpected frequency-dependent excitatory feedback loop between pyramidal cell activity and dendritic spiking that is mediated by NMDAR co-agonists and astrocytes. Importantly, disrupting this feedback loop at the level of astrocytes impaired spatial memory.

## Results

### Control of dendritic spiking by the NMDAR co-agonist D-serine

In a first set of experiments, we investigated dendritic spiking of CA1 pyramidal cells in acute hippocampal slices by iontophoretic glutamate application onto their dendrites in the stratum radiatum (Müller et al. 2012) (**Fig. 1a-b**). Increasing synaptic input was emulated by increasing the iontophoretic current and thus the amount of ejected glutamate. As expected, the amplitude of the somatically recorded voltage response increased (**Fig. 1c**, black traces and dots) until a threshold was passed and dendritic spikes composed of a fast and sodium channel-dependent component and a slow component (Ariav et al. 2003; Losonczy and Magee 2006) were detected (**Fig. 1c**, orange dots and traces). Throughout this study, we used the threshold stimulus that just evoked a dendritic spike and the amplitude of the slow component as parameters describing dendritic spikes. We then used acute inhibition of NMDARs by APV to establish the extent to which NMDARs control dendritic spikes. APV increased the threshold stimulus and decreased the slow component of the dendritic spikes significantly (**Fig. 1d**) reflecting the well-established NMDAR-dependence of these dendritic spikes (Ariav et al. 2003; Harnett et al. 2012; Losonczy and Magee 2006; Schiller et al. 2000). We next tested if increasing the occupancy of NMDAR co-agonist binding site by application of exogeneous D-serine (10 μM) alters dendritic spiking and found the opposite effect compared to NDMAR blockade (**Fig. 1e**). This indicates that the NMDAR co-agonist binding site is not saturated in our experimental condition and that its occupancy dynamically regulates dendritic spiking. Control experiments without drug application revealed that both stimulus threshold and the amplitude of the slow component were stable over time (not illustrated; baseline and after ~ 10 minutes; threshold: 0.56 ± 0.08 μA and 0.56 ± 0.08 μA, t(5) = 0.31, n = 6, p = 0.76, paired Student’s t-test; slow component: 13.22 ± 2.74 mV and 13.39 ± 2.69 mV, n = 6, t(5) = 0.51, p = 0.63, paired Student’s t-test). In addition, metabotropic glutamate receptors were also not involved (**Suppl. Fig. 1**). Please note that here and throughout the study we used acute experimental manipulations whenever possible because absolute thresholds and amplitudes depend on the exact experimental configuration (e.g. iontophoretic pipettes, distance of stimulation site from cell body).

**Figure 1:**
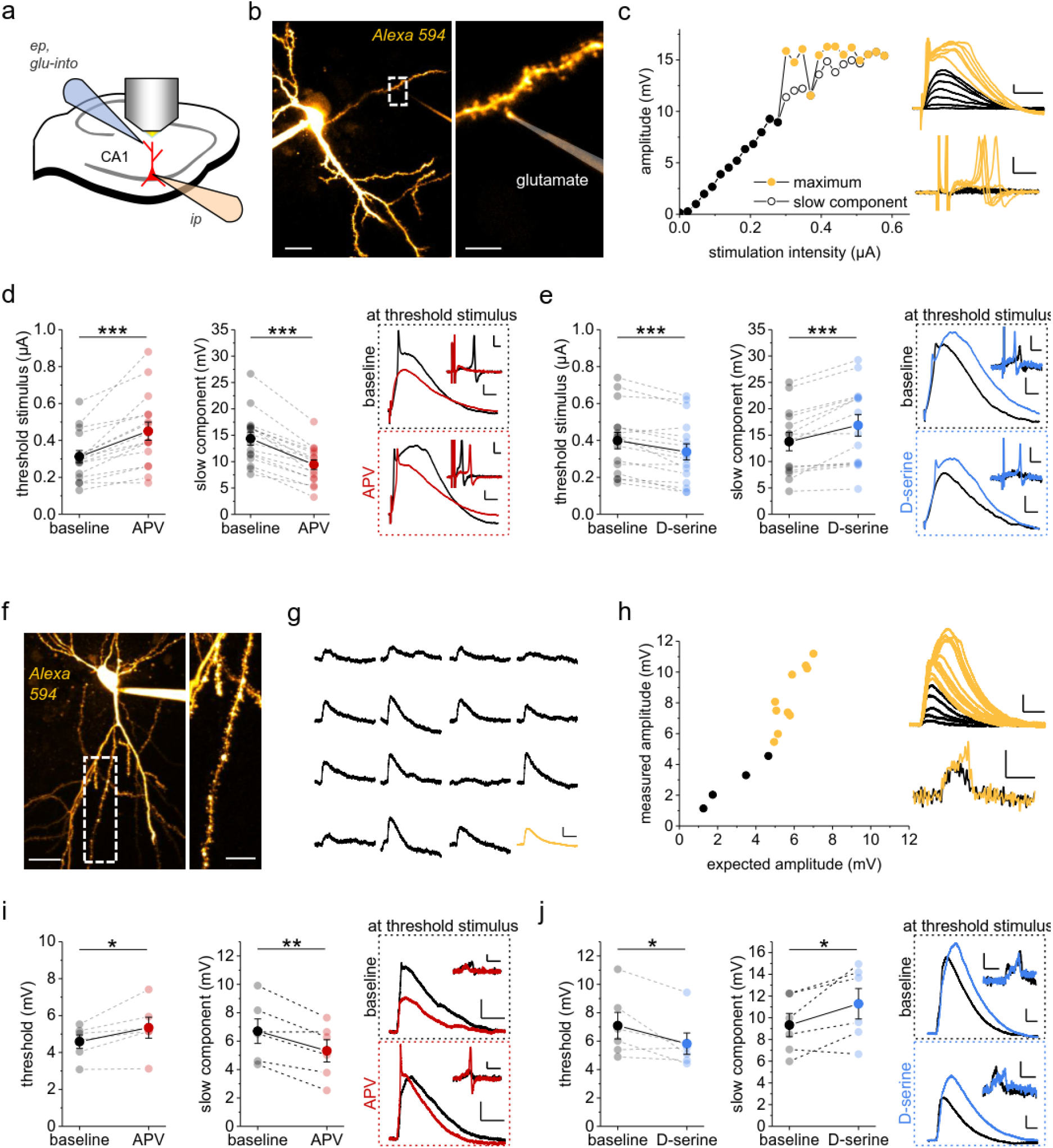
Threshold and amplitude of dendritic spikes are controlled by N-methyl-D-aspartate receptor (NMDAR) co-agonist levels. **a)** Schematic of experiments. Whole-cell patch clamp recordings (ip, intracellular pipette, current clamp) from CA1 pyramidal cells (red) combined with iontophoretic application of glutamate (ep, extracellular pipette) and two-photon excitation (2PE) fluorescence microscopy. Dyes were added to visualize the patched cell and the iontophoresis pipette (Alexa Fluor 594, 40 μM and 50 μM, respectively). **b)** Sample images of a typical recording (left panel: scale bar 20 μm; right panel: zoomed in on dendrite, dashed box in left panel, scale bar 5 μm). **c)** Sample experiment. Dependence of the somatically recorded depolarizations on the iontophoretic current (left panel, filled circles: maximum amplitude, black below dendritic spike threshold and yellow above, empty circles: amplitude of slow component). Right panels: sample traces corresponding to left panel. Upper right panel: voltage, scale bars 2 mV and 20 ms. Lower right panel: dV/dt, scale bars 5 mV/ms and 2 ms. Note the difference of time scales and that the first two deflections represent the dV/dt of the stimulus artifact. **d)** NMDAR blockade by D-APV (50 μM) increased the threshold stimulus (left panel, smallest iontophoretic current that elicited a dendritic spike, 0.31 ± 0.03 μA vs. 0.45 ± 0.05 μA, n = 16, p < 0.0001) and reduced the slow component (middle panel, 14.36 ± 1.23 mV vs. 9.42 ± 0.90 mV, n = 16, p < 0.0001). Right top panel: sample traces recorded with the baseline threshold stimulus. Right bottom panel: sample traces recorded with the threshold stimulus in D-APV (baseline: black traces; D-APV: red traces; scale bars 2 mV and 10 ms; insets: dV/dt, scale bars 5 mV/ms and 2 ms). **e)** D-serine (10 μM) significantly decreased the threshold stimulus (left panel, 0.40 ± 0.04 μA vs. 0.34 ± 0.04 μA, n = 16, p = 0.00011) and increased the amplitude of the slow component (middle panel, 13. 77 ± 1.73 mV vs. 16.87 ± 2.05 mV, n = 14, p = 0.00024). Right panels: sample traces of somatic voltage and dV/dt (insets) as before (upper traces with threshold stimulus during baseline; lower traces with threshold stimulus in D-serine, scale bars for voltage 2 mV and 10 ms and for insets 5 mV/ms and 2 ms). **f)** Two-photon glutamate uncaging experiments. Example of a CA1 pyramidal cell filled with Alexa Fluor 594 (100 μM, left panel, scale bar 40 μm). Right panel: investigated dendrite with uncaging locations at dendritic spines (scale bar 10 μm). **g)** Examples of uncaging-evoked somatic EPSPs (uEPSP, scale bars 0.5 mV and 20 ms, average uEPSP in orange). **h)** Left panel: comparison of the measured amplitudes of somatic responses evoked by quasi-simultaneous stimulation of a set of spines with the sum of single-spine uEPSPs of the same set (expected amplitude for summation, black: no spikes, orange: spikes detected). Right top panel: individual responses corresponding to the left panel (scale bars 2 mV and 20 ms). Right bottom panel: dV/dt of traces above (scale bars 1 mV/ms and 5 ms). **i)** Left panel: thresholds of dendritic spikes were significantly increased with blocked NMDARs (D-APV, 40 μM, red, 4.59 ± 0.37 mV vs. 5.34 ± 0.57 mV, n = 6, p = 0.039). Middle panel: NMDAR inhibition decreased the slow component of the dendritic spike (6.70 ± 0.86 mV vs. 5.32 ± 0.78 mV, n = 6, p = 0.0097). Right panels: sample traces as in **d** (scale bars 2 mV and 50 ms, inset dV/dt scale bars 1 mV/ms and 10 ms). **j)** Application of saturating concentrations of D-serine (10 μM, blue) decreased the dendritic spike threshold (left panel, 7.09 ± 0.94 mV vs. 5.82 ± 0.75 mV, n = 6, p = 0.049) and increased the slow component of the dendritic spike (middle panel, 9.33 ± 1.07 mV vs. 11.29 ± 1.40 mV, n = 6, p = 0.045). Right panels: example traces as in **e** (scale bars 2 mV and 20 ms, inset dV/dt scale bars 1 mV/ms and 10 ms). Paired Student’s t-tests throughout.

The bi-directional modification of dendritic spiking by NMDAR blockade and co-agonist site saturation were then confirmed using two-photon glutamate uncaging for stimulating an increasing number of dendritic spines (Losonczy and Magee 2006; Remy et al. 2009) (**Fig. 1f-h**). Again, inhibition of NMDARs by APV decreased the amplitude of the slow spike component whereas exogenous D-serine had the opposite effect (**Fig. 1i-j**). For these uncaging experiments, the spike threshold was calculated as the expected excitatory postsynaptic potential (EPSP) amplitude at which dendritic spikes first occurred, i.e., as the arithmetic sum of the amplitudes of single spine uncaging-evoked EPSPs (uEPSPs) that was sufficient to trigger a dendritic spike. Similar to iontophoretic stimulation, NMDAR inhibition increased whereas D-serine application decreased the threshold (**Fig. 1i-j**). These observations were not due to changes of AMPAR-mediated synaptic transmission because neither iontophoretically evoked miniature-like EPSPs nor uEPSPs elicited by glutamate uncaging were affected by any of the drugs (p > 0.10 throughout, paired Student’s t-tests).

Our observations are in line with previous studies showing that NMDARs contribute to the late phase of dendritic spikes (Ariav et al. 2003; Losonczy and Magee 2006). Similarly, blockade of NMDARs was previously shown to increase the dendritic spike threshold to a variable degree (Ariav et al. 2003; Losonczy and Magee 2006) and increasing NMDAR co-agonist levels decreased the spike threshold in our experiments. Therefore, our results indicate that boosting NMDAR function and thus NMDAR-mediated depolarization via co-agonist supply could facilitate the activation of voltage-dependent ion channels, for example sodium channels mediating the initial fast component of dendritic spikes (Ariav et al. 2003; Losonczy and Magee 2006).

We and others have previously shown that astrocytic Ca^2+^ signaling can regulate the supply of D-serine to NMDARs (Henneberger et al. 2010; Rasooli-Nejad et al. 2014; Robin et al. 2018; Takata et al. 2011). Therefore, triggering astrocytic Ca^2+^ changes should affect dendritic spikes. A potent trigger of astrocytic Ca^2+^ signals is activation of their endocannabinoid receptors (CBRs) (Navarrete and Araque 2008; Rasooli-Nejad et al. 2014; Robin et al. 2018). After verifying that activation of CBR signaling using WIN55 induced astrocytic Ca^2+^ signals (**Fig. 2a-b**), we tested if this manipulation also affects dendritic spiking. Indeed, we found that application of WIN55 lowered the stimulus threshold and increased the slow component of dendritic spikes (**Fig. 2c**). Importantly, we did not observe this when the experiments were performed in the presence of exogenous D-serine to saturate the NMDAR co-agonist binding site (**Fig. 2d**). This finding indicates that activation of CBR induces astrocytic Ca^2+^ signals and modifies dendritic spiking via the NMDAR co-agonist binding site.

**Figure 2:**
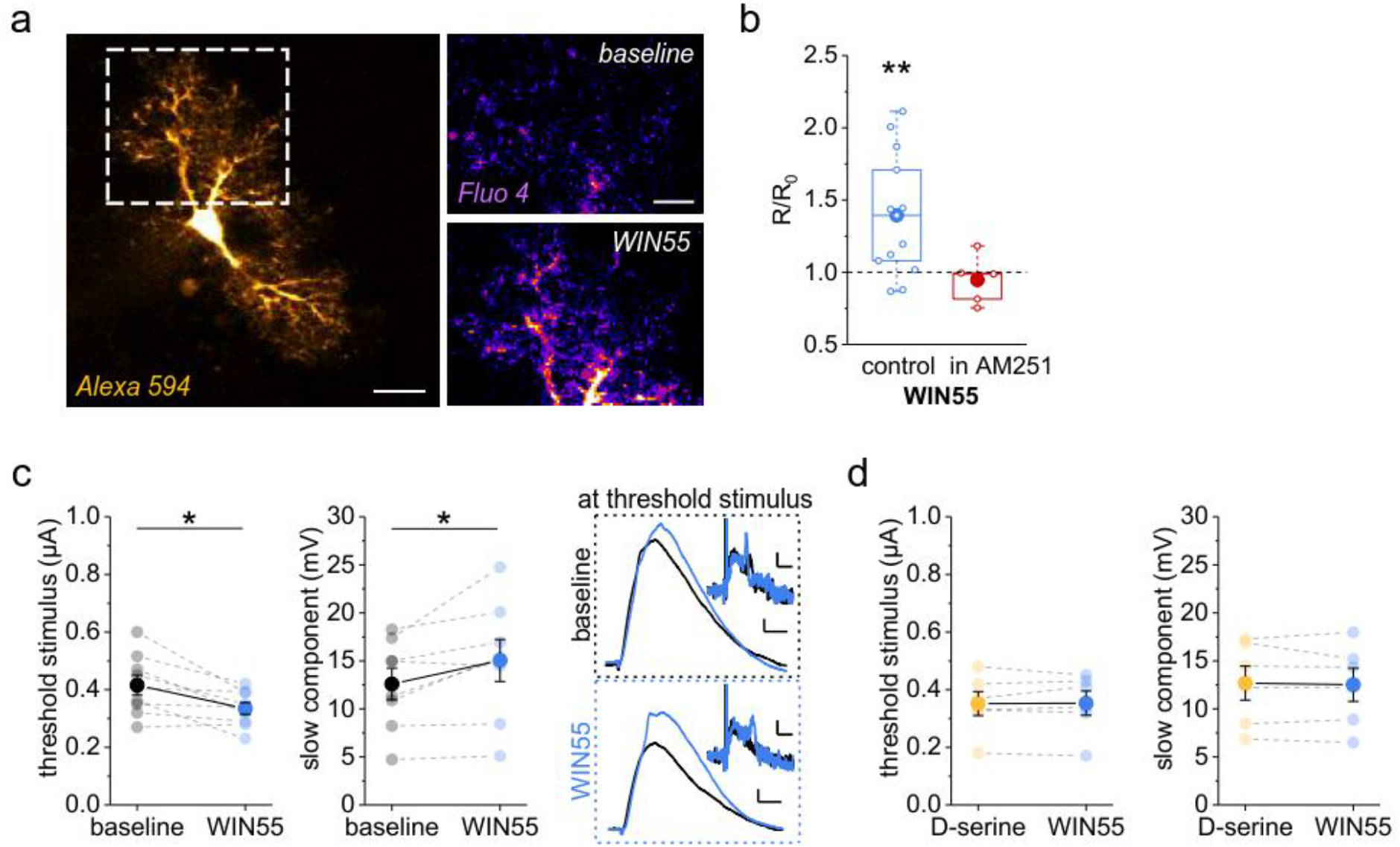
Endocannabinoid receptor (CBR) activity controls astrocytic Ca^2+^ and dendritic spike threshold and amplitude. **a)** Example astrocyte filled with the Ca^2+^-sensitive dye Fluo-4 (200-400 μM) and the Ca^2+^-insensitive dye Alexa Fluor 594 (40 μM, left panel, scale bar 10 μm, whole-cell patch clamp). Right upper panel: baseline Fluo-4 fluorescence intensity. Right lower panel: Fluo-4 intensity during application of the CBR agonist WIN55 (WIN55,212-2, 10 μM). Both right panels: average of 10 frames, scale bar 5 μm. **b)** Change of the fluorescence intensity ratio (R, Fluo-4 / Alexa 594) quantified as R in WIN55 relative to baseline before WIN55 application (R_0_, also see Methods). Significant increase by WIN55 (blue, 1.39 ± 0.12, n = 13, p = 0.005, one-population Student’s t-test) but not in the presence of the CBR inverse agonist AM251 (5 μM, red, 0.95 ± 0.08, n = 5, p = 0.52, one-population Student’s t-test). **c)** CBR activation by WIN55 (blue, 1 μM) decreased the threshold stimulus of dendritic spikes (left panel, 0.42 ± 0.03 μA vs. 0.33 ± 0.02 μA, n = 9, p = 0.012, paired Student’s t-test) and increased the amplitude of the slow dendritic spike component (middle panel: 12.59 ± 1.65 mV vs. 15.04 ± 2.18 mV, n = 8, p = 0.034, paired Student’s t-test). Right panels: sample EPSP traces (scale bars: 2 mV and 10 ms) and dV/dt (insets, scale bars: 1 mV/ms and 5 ms) at the baseline threshold stimulus (upper traces) and at the threshold stimulus in WIN55 (lower traces). Both panels black for baseline and blue for WIN55. **d)** In presence of saturation concentrations of D-serine (10 μM) WIN55 fails to change the threshold stimulus (left panel, 0.35 ± 0.04 μA vs. 0.35 ± 0.04 μA, n = 6, p = 0.87, paired Student’s t-test) and the slow dendritic spike component (right panel, 12.69 ± 1.77 mV vs. 12.52 ± 1.73 mV, n = 6, p = 0.64, paired Student’s t-test).

### Pyramidal cell activity controls dendritic spiking via the NMDAR co-agonist site

The observations above were obtained with pharmacological activation of astrocytic CBRs. We next explored if endogenous CBR activation would lead to similar effects on dendritic spiking. Postsynaptic depolarization is a powerful trigger of endocannabinoid release (Di Marzo et al. 1994; Stella et al. 1997). We therefore induced postsynaptic depolarization of CA1 pyramidal cells by antidromic stimulation of their axons in the alveus (**Fig. 3a**, **Suppl. Fig. 2**), which reliably activated ~ 40% of pyramidal cells and over a broad range of stimulation frequencies (**Suppl. Fig. 2e-j**). We then investigated if this stimulation protocol induces astrocytic activity via endocannabinoid signaling. Indeed, antidromic activation of CA1 pyramidal cells evoked astrocytic Ca^2+^ signals, which were abolished by CBR inhibition using AM251 (**Fig. 3a-c**).

**Figure 3:**
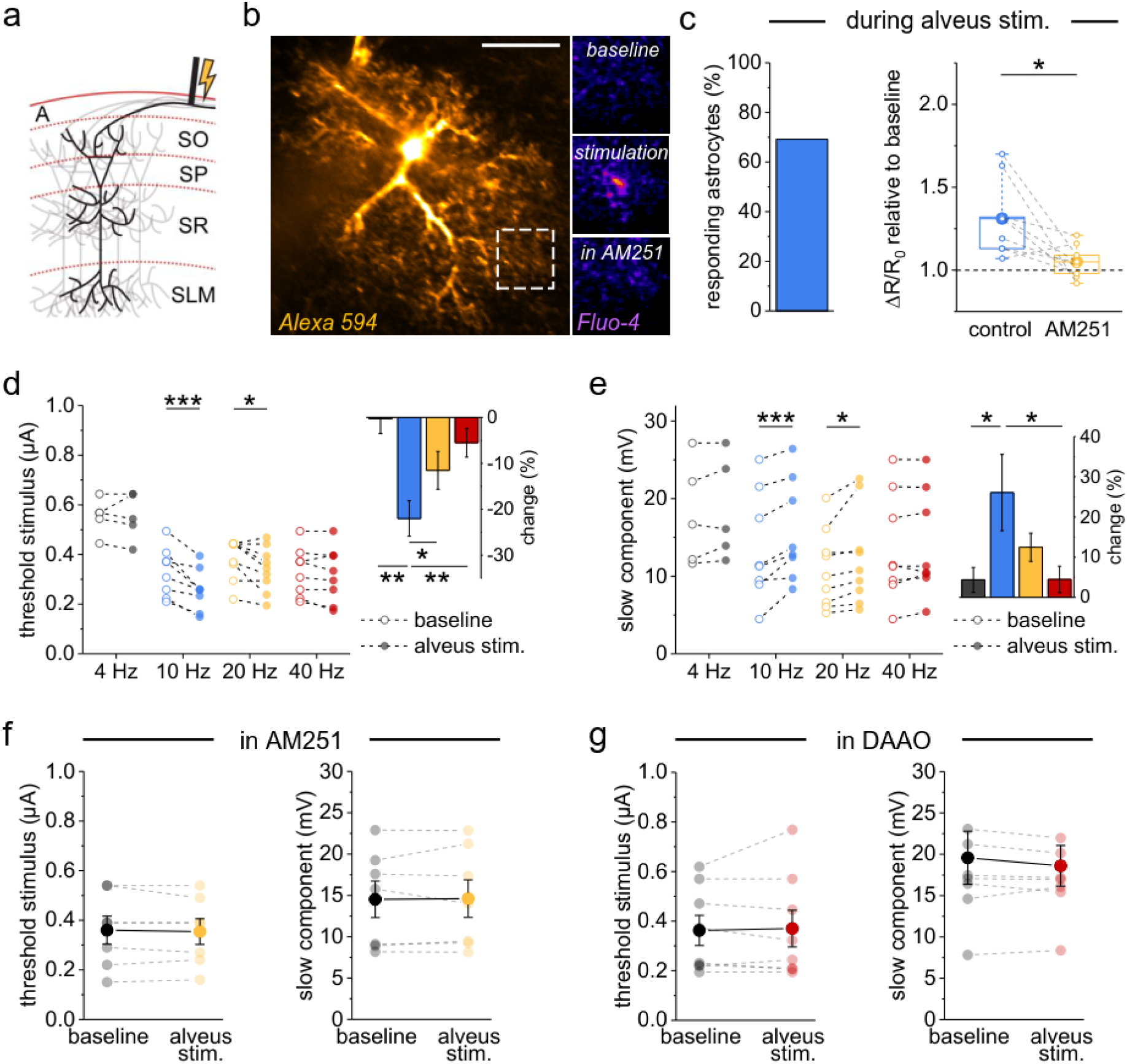
Pyramidal cell activity increases astrocytic Ca^2+^ signaling and promotes dendritic spiking via CBRs and NMDAR co-agonist supply. **a)** Stimulation of axons in the alveus (A) retrogradely increases the activity of CA1 pyramidal cells (SO, stratum oriens; SP, stratum pyramidale; SR, stratum radiatum; SLM, stratum lacunosum moleculare). **b)** Example astrocyte (whole-cell patch clamp, 40 μM Alexa Fluor 594 and 400 μM Fluo-4, scale bar: 20 μm). Left panel: Alexa Fluor 594. Right panels: Fluo-4, average of 20 frames during baseline, alveus stimulation and alveus stimulation in the presence of the CBR inverse agonists AM251 (5 μM), from top to bottom. Quantification of Ca^2+^ signals by the fluorescence intensity ratio (R, Fluo-4 / Alexa). Its overall changes (ΔR) relative to its resting value (R_0_) during alveus stimulation were compared to the baseline period before stimulation (also see Methods). **c)** 69.2% of the astrocytes showed an increase of at least 5% (left panel). In the responders, the increase was blocked by AM251 (5 μM; 1.31 ± 0.07 vs 1.05 ± 0.03, n = 9, p = 0.016, paired Student’s t-test). **d)** Left panel: the threshold stimulus of dendritic spikes was decreased by alveus stimulation at 10 Hz (0.33 ± 0.03 μA vs. 0.26 ± 0.03 μA, n = 8, p < 0.0001, repeated measures ANOVA and post-hoc Fisher’s LSD) and 20 Hz (0.38 ± 0.03 μA vs. 0.34 ± 0.03 μA, n = 9, p = 0.031, Friedman test and post-hoc Wilcoxon signed-rank test) but not 4 Hz (0.55 ± 0.03 μA vs. 0.55 ± 0.04 μA, n = 5, p = 1, paired Student’s t-test, separate set of experiments) and 40 Hz (0.33 ± 0.03 μA vs. 0.32 ± 0.04 μA, n = 8, p = 0.31, repeated measures ANOVA and post-hoc Fisher’s LSD). Open circles: baseline. Filled circles: alveus stimulation. Right panel: change relative to baseline (one-way ANOVA with post-hoc Fisher’s LSD). **e)** Left panel: changes of the slow component amplitude displayed a similar frequency dependence (10 Hz: 13.70 ± 2.47 mV vs. 15.74 ± 2.29 mV, n = 8, p < 0.0001, repeated measures ANOVA with post-hoc Fisher’s LSD; 20 Hz: 9.75 ± 1.37 mV vs. 11.08 ± 1.81 mV, n = 8, p = 0.022, repeated measures ANOVA with post-hoc Fisher’s LSD; 4 Hz: 17.97 ± 2.98 mV vs. 18.61 ± 2.94 mV, n = 5, p = 0.23, paired Student’s t-test, separate set of experiments; 40 Hz: 13.70 ± 2.47 mV vs. 14.00 ± 2.39 mV, n = 8, p = 0.41, repeated measures ANOVA with post-hoc Fisher’s LSD). Open circles: baseline. Filled circles: alveus stimulation. Right panel: change relative to baseline (Kruskal-Wallis test with post-hoc Mann-Whitney tests). **f)** In the presence of AM251 (5 μM, orange) alveus stimulation did not affect the dendritic spike threshold stimulus (left panel, 0.36 ± 0.06 μA vs. 0.35 ± 0.05 μA, n = 7, p = 0.54, paired Student’s t-test) or its slow component amplitude (right panel, 14.53 ± 2.21 mV vs. 14.61 ± 2.27 mV, n = 7, p = 0.85, paired Student’s t-test). **g)** Presence of the enzyme D-amino acid oxidase (DAOO, 0.17 U/ml, red) prevented the change of the dendritic spike threshold stimulus (0.36 ± 0.06 μA vs. 0.37 ± 0.07 μA, n = 8, p = 0.73, paired Student’s t-test) and slow component amplitude (19.58 ± 3.21 mV vs. 18.60 ± 2.46 mV, n = 8, p = 0.27, paired Student’s t-test). Throughout panels * p < 0.05, ** p < 0.01, *** p < 0.001.

According to our previous results, such stimulation of pyramidal cell and astrocytic activity should increase the extracellular NMDAR co-agonist level and thereby affect dendritic spiking. We tested this prediction by monitoring dendritic integration before and during alveus stimulation at various frequencies and found that stimulation at 10 and 20 Hz but not at 4 and 40 Hz reduced the stimulus threshold of dendritic spikes (**Fig. 3d**) and increased the slow component of dendritic spikes (**Fig. 3e**). This reveals that population activity of pyramidal cells facilitates the generation of dendritic spikes in a frequency-dependent manner. Similar experiments in the presence of the CBR antagonist AM251 failed to change dendritic spiking and thus demonstrate that CBRs mediate this positive, frequency-dependent feedback loop (alveus stimulation at 20 Hz, **Fig. 3f**). A change of dendritic spiking was also not observed when we performed this experiment in the presence of D-serine (not illustrated; alveus stimulation at 20 Hz, threshold: 0.44 ± 0.05 μA and 0.45 ± 0.05 μA, n = 6, t(5) = 1, p = 0.36, paired Student’s t-test; slow component: 20.34 ± 2.54 mV and 20.05 ± 2.46 mV, n = 6, t(5) = 0.55, p = 0.61, paired Student’s t-test). We next asked if indeed D-serine is the endogenous co-agonist involved. The identity of the relevant co-agonist can be established by incubating brain slices with D-amino acid oxidase (DAAO) to degrade endogenous D-serine (Mothet et al. 2000; Panatier et al. 2006; Papouin et al. 2012). Repeating our experiments in the presence of DAAO indeed prevented a modulation of dendritic spiking by alveus stimulation (10 Hz), which reveals that indeed D-serine is involved (**Fig. 3g**). Consistent with a transient release of D-serine triggered by pyramidal cell activity, dendritic spiking returned to its baseline within five minutes after alveus stimulation (**Suppl. Fig. 3**).

### Mechanism underlying the frequency dependence of dendritic spiking modulation

Next, we dissected the cellular mechanisms underlying the unexpected frequency dependence of activity-induced changes of dendritic integration. While it may not be surprising that low frequency stimulation is insufficient to activate this pathway, the absence of changes of dendritic integration at 40 Hz was less expected. We had already established that pyramidal cells activated by alveus stimulation reliably follow stimulation frequencies between 4 and 40 Hz (**Suppl. Fig. 2h-j**). Thus, the frequency dependence could be caused at the level of CBR-dependent astrocyte Ca^2+^ signaling or co-agonist release or at the level of excitability of pyramidal cell dendrites, the presumed location of endocannabinoid release. If the latter is the case, then astrocytic Ca^2+^ signals are expected to also display a frequency-dependent modulation by alveus stimulation. Indeed, alveus stimulation at 10 Hz was more efficiently inducing astrocytic Ca^2+^ transients than at 40 Hz (**Fig. 4a-c**, **Suppl. Fig. 4**), which again could arise from frequency-dependent astrocytic Ca^2+^ signaling or pyramidal cell endocannabinoid release. Because a frequency-dependence of pyramidal cell excitability can be conferred by HCN channels to pyramidal cells and especially to their dendrites (Hu et al. 2002; Narayanan and Johnston 2007), we next tested their involvement by performing experiments in the presence of the HCN inhibitor ZD7288. In these experiments, we first established the whole-cell patch clamp configuration, washed in ZD7288 and compensated the hyperpolarization of the recorded pyramidal cell due to HCN blockade (−4.99 ± 1.06 mV, n = 18) by a constant current injection before testing dendritic spiking. Indeed, we found that alveus stimulation at 10 Hz was not affecting the stimulus threshold and slow components of dendritic spikes any longer in the presence of ZD7288 (**Fig. 4d-e**, orange). However, direct activation of CBRs by WIN55 was still effective (**Fig. 4d-e**, blue) indicating that the frequency dependence is likely caused by dendritic mechanisms that involve HCN channels.

**Figure 4:**
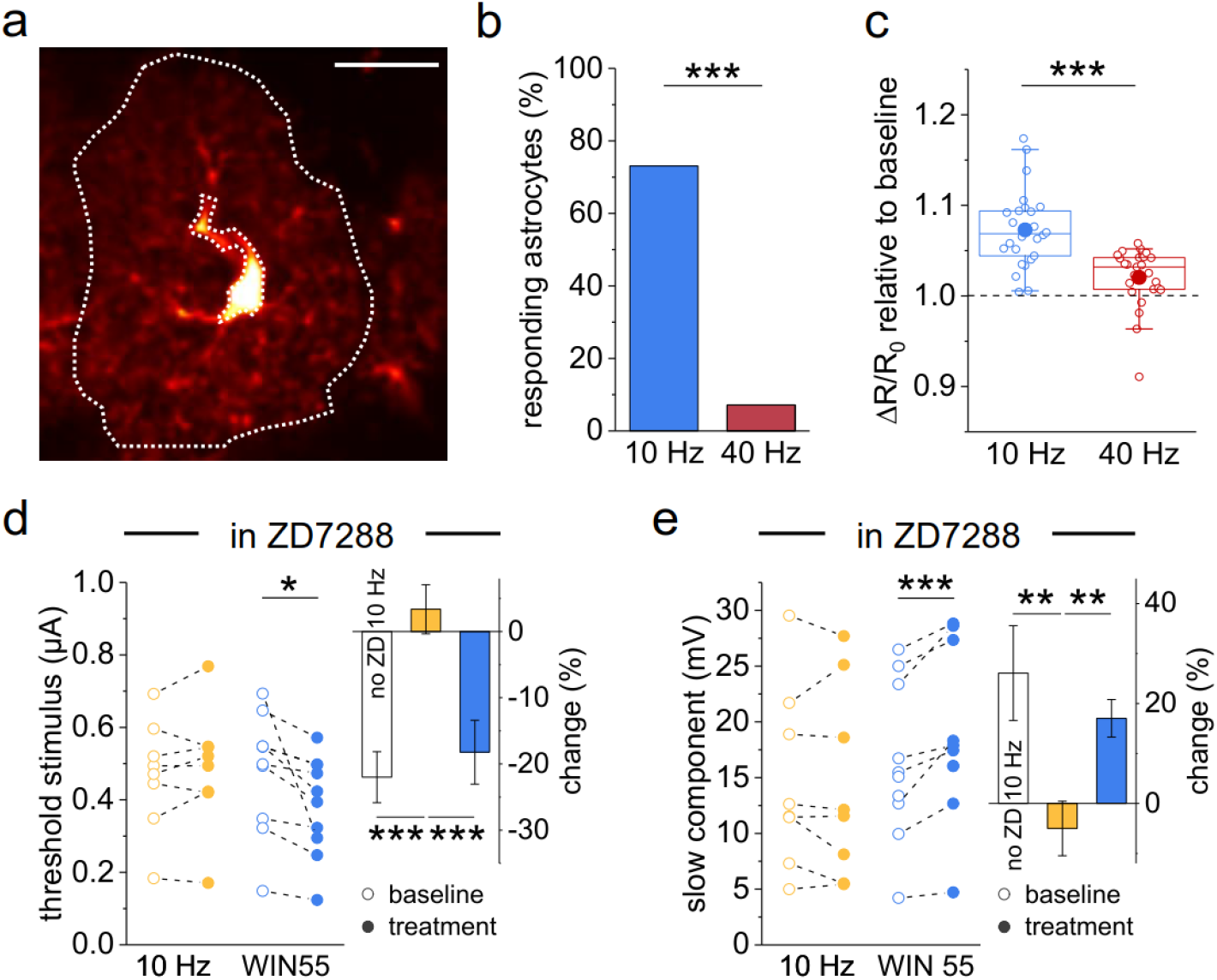
Mechanisms underlying the frequency-dependence of dendritic spike modulation by alveus stimulation. **a)** Representative astrocyte expressing GCaMP5g (not illustrated) and tdTomato (red, scale bar: 20 μm; region of interest: dotted while line, sparing the cell body and major branches). Quantification of Ca^2+^ signals by the fluorescence intensity ratio (R, GCaMP5g / tdTomato) and its changes (ΔR) over its resting value (R_0_). Comparison of ΔR/R0 during alveus stimulation with the baseline period before stimulation (also see Methods). **b)** Percentage of astrocytes with an increase of Ca^2+^ signaling increase of > 5% was significantly larger with 10 Hz (blue) stimulation than with 40 Hz (73.1 % vs. 7.14 %, Fisher’s exact test, p < 0.0001). **c)** Change of ΔR/R0 during alveus stimulation across all cells (10 Hz vs 40 Hz: 1.07 ± 0.008 vs. 1.02 ± 0.007, n = 26/25, p < 0.0001, Mann-Whitney test). **d-e)** Inhibition of hyperpolarization-activated cyclic nucleotide-gated (HCN) channels using ZD7288 (10 μM) prevented the modulation of dendritic spikes thresholds and slow component amplitudes by alveus stimulation while the effect of direct CBR stimulation by WIN55 (1 μM) was preserved. **d)** Left panel: dendritic spike threshold stimulus, 10 Hz (0.47 ± 0.05 μA vs. 0.49 ± 0.06 μA, n = 8, p = 0.33, paired Student’s t-test), WIN55 (0.48 ± 0.05 μA vs. 0.38 ± 0.04 μA, n = 10, p = 0.025, paired Student’s t-test). Right panel: change during treatment (solid circles, left panel) relative to baseline (open circles, left panel), Kruskal-Wallis test with post-hoc Mann-Whitney test. **e)** Left panel: dendritic spike slow component amplitude, 10 Hz (14.75 ± 2.87 mV vs. 14.27 ± 3.05 mV, n = 8, p = 0.52, paired Student’s t-test), WIN55 (16.24 ± 2.21 mV vs. 18.79 ± 2.41 mV, n = 10, p = 0.0010, paired Student’s t-test). Right panel: change during treatment (solid circles, left panel) relative to baseline (open circles, left panel), Kruskal-Wallis test with post-hoc Mann-Whitney test. Throughout panels * p < 0.05, ** p < 0.01, *** p < 0.001. In the right panels of d and e, the white control bar with 10 Hz alveus stimulation represents experiments without ZD7288 from **Fig. 3d-e**.

### Astrocytic type 1 endocannabinoid receptors (CB1Rs) in dendritic spiking and place memory

Our results indicate that astrocytic CBRs close a positive feedback loop between pyramidal cell activity and their dendritic spiking via NMDAR co-agonist signaling. To directly establish astrocytic CBR involvement, we deleted astrocytic CB1Rs by injecting a transgenic mouse line GLASTcreERT2 (Mori et al. 2006) crossed with the CB1R fl/fl line (Marsicano et al. 2003) and the reporter line (flox-stop-tdTomato) (Madisen et al. 2010) with tamoxifen (aCB1KO). We used sham injected littermates (sham) and wildtype animals (WT) as controls (**Fig. 5**). We first confirmed that tdTomato expression was near-exclusively restricted to astrocytes in the hippocampal CA1 region (**Suppl. Fig. 5**). In addition, no major differences of basic properties of CA3-CA1 synaptic transmission were detected between slices from these mice (**Suppl. Fig. 6**, **Supp. Table 1**). If astrocytic CB1Rs mediate frequency-dependent changes of dendritic spiking, then their removal in aCB1KO mice should disrupt this positive feedback. We therefore tested dendritic spiking evoked by iontophoretic glutamate application in the territory of a tdTomato-expressing astrocyte before and during alveus stimulation (**Fig. 5a-b**). Indeed, we found that alveus stimulation at 10 Hz was unable to affect the threshold stimulus and the amplitude of the slow component of dendritic spikes, while this effect was however preserved in slices from WT and sham animals (**Fig. 5c-d**). This indicates that astrocytes CB1Rs mediate the positive and frequency-dependent feedback loop between pyramidal cell activity and their dendritic integration (see **Suppl. Fig. 9** for a schematic).

**Figure 5:**
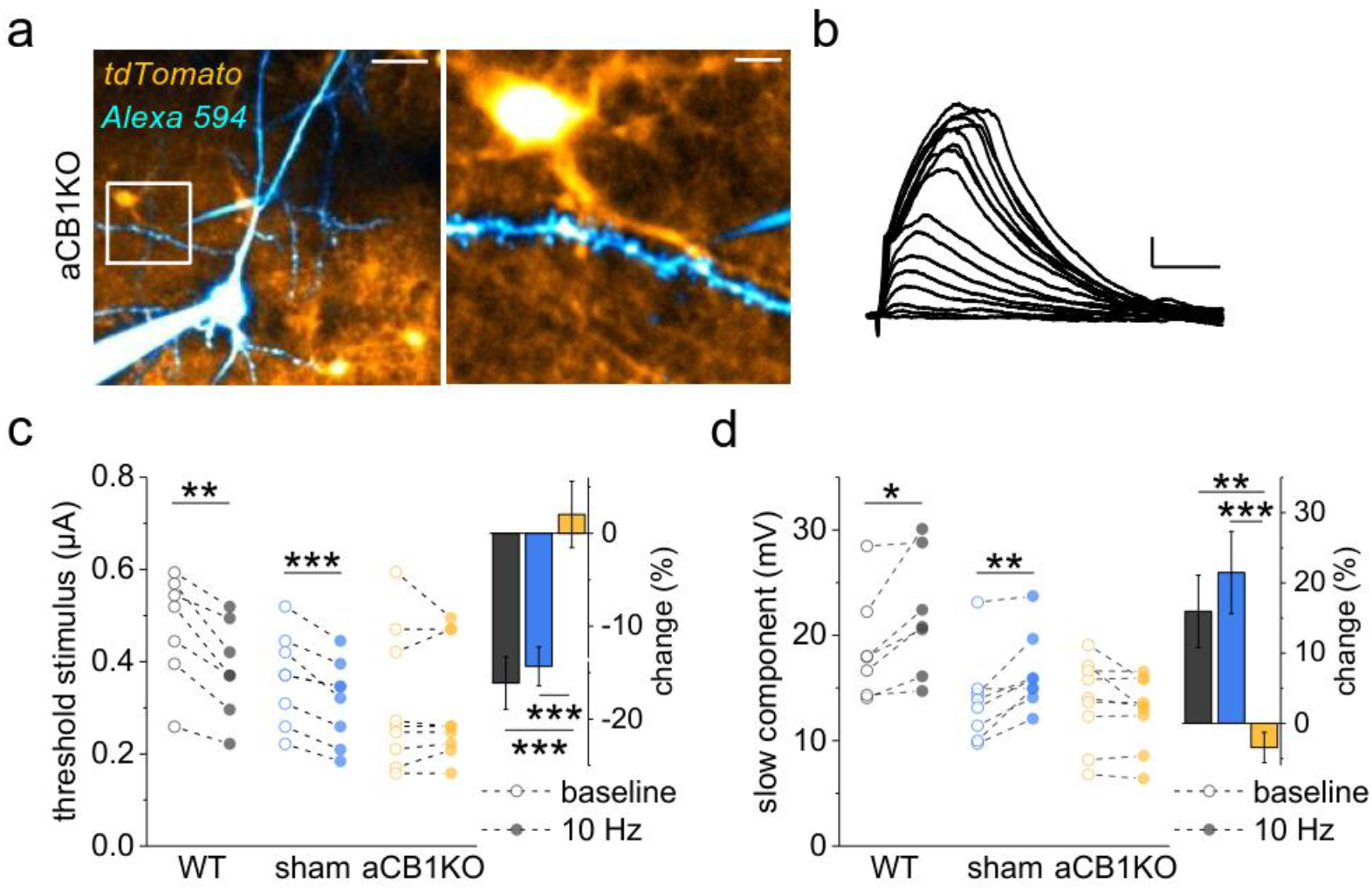
Deletion of astrocytic type 1 CBRs (CB1Rs) disrupts the frequency-dependent modulation of dendritic spiking by pyramidal cell activity. **a)** Example of a CA1 pyramidal cell filled with Alexa Fluor 594 (40 μM, blue, whole-cell patch clamp), an iontophoresis pipette (blue) and expression of tdTomato as an indicator of conditional deletion of CB1Rs in astrocytes (aCB1KO). Left panel: scale bar 20 μm. Right panel: enlarged region outlined by white box in right panel, scale bar 5 μm. **b)** Sample EPSPs recorded in aCB1KO mouse in response to increasing iontophoretic glutamate applications (scale bar: 3 mV and 20 ms). **c)** dendritic spikes recorded in aCB1KO mice lack the stimulus threshold decrease during 10 Hz alveus stimulation observed in wildtype (WT) and sham control mice (left panel; WT: 0.47 ± 0.04 μA vs. 0.38 ± 0.04 μA, n = 7, p = 0.0017, paired Student’s t-test; sham: 0.36 ± 0.03 μA vs. 0.31 ± 0.03 μA, n = 8, p = 0.00071, paired Student’s t-test; aCB1RKO: 0.31 ± 0.05 μA vs. 0.31 ± 0.04 μA, n = 9, p = 1.00, Wilcoxon signed-rank test). Right panel: change during 10 Hz alveus stimulation (filled circles, left panel) relative to baseline (open circles, left panel). One-way ANOVA with post-hoc Fisher’s LSD. **d)** Slow component amplitudes of dendritic spikes recorded in aCB1KO mice lack the stimulus threshold increase during 10 Hz alveus stimulation observed in WT and sham mice (WT: 18.81 ± 1.91 mV vs. 21.93 ± 2.20 mV, n = 7, p = 0.021, paired Student’s t-test; sham: 13.83 ± 1.50 mV vs. 16.41 ± 1.29 mV, n = 8, p = 0.0056, paired Student’s t-test; aCB1KO: 13.73 ± 1.36 mV vs. 12.83 ± 1.14 mV, n = 9, p = 0.14, paired Student’s t-test). Right panel: change during 10 Hz alveus stimulation (filled circles, left panel) relative to baseline (open circles, left panel). One-way ANOVA with post-hoc Fisher’s LSD. Throughout panels * p < 0.05, ** p < 0.01, *** p < 0.001.

Because NMDAR-driven dendritic spikes in CA1 pyramidal cells have been implicated in the formation of place memory (for review Sheffield and Dombeck 2019), we next asked if genetic deletion of astrocytic CB1Rs impairs behaviors that require the encoding of a location. First, we tested object location memory by allowing animals to explore two objects in a first session (acquisition) and then moving one of the objects to a new location (recall, **Fig. 6a**). An increased exploration of the displaced object during the recall trial indicates that the animal has encoded the initial location and is able to discriminate the displaced object. Sham injected and aCB1KO animals showed neither an initial preference for object locations during acquisition (0.08 ± 0.03 vs. −0.002 ± 0.07, n = 10/12, t(20) = 0.98, p = 0.34, Student’s t-test), nor did the overall exploration of the objects differ on the test day (recall, **Fig. 6b**). We also did not detect differences between the groups regarding the exploration of the test arena (**Suppl. Fig. 7a-d**) and the spontaneous alternation in the Y-maze, indicating intact spatial working memory (**Suppl. Fig. 7e-h**). In contrast, aCB1KO mice did not display a preference for the displaced object in the recall trial compared to sham injected animals (**Fig. 6c**), indicating impaired object location memory.

**Figure 6:**
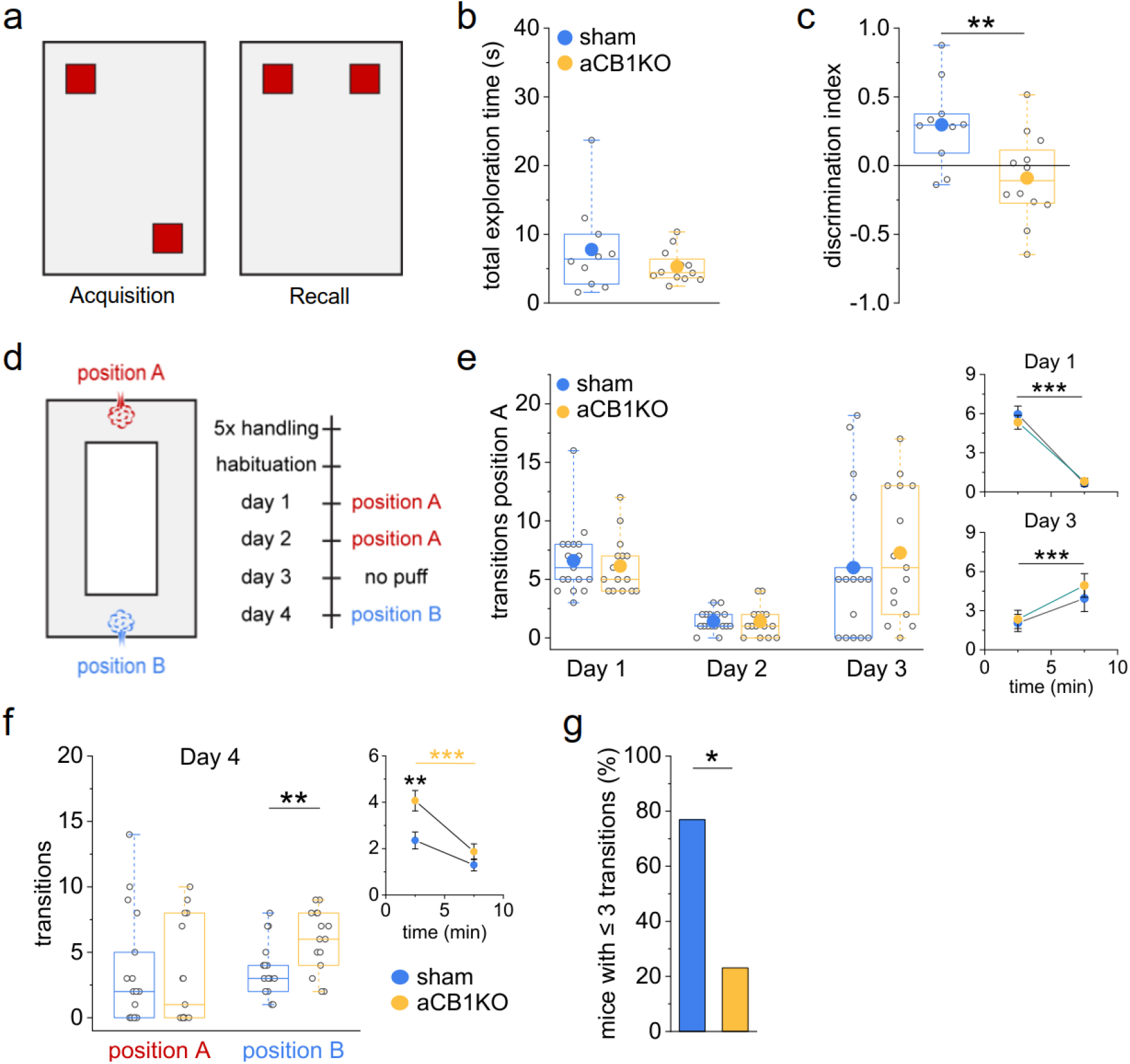
Disruption of astrocytic CB1R-mediated signaling impairs formation of spatial memory. **a)** Schematic arena for testing object location memory. Two identical objects (red squares) are explored by the animals (10 min) during acquisition. During testing (Recall), one object is moved to a new location and mice are allowed for freely explore the objects (5 min). **b)** Total exploration time did not differ between sham control animals and animals with conditional deletion of CB1Rs in astrocytes (aCB1KO) (7.78 ± 2.07 s vs. 5.30 ± 0.69 s, n = 10/12, p = 0.54, Mann-Whitney test). **c)** Sham control mice explored the object in the novel location significantly more whereas aCB1KO mice did not discriminate (0.30 ± 0.10 vs. −0.09 ± 0.09, n = 10/12, p = 0.0098, Student’s t-test). **d)** Schematic and timeline of passive place avoidance test with two possible positions for a triggered air puff when animals entered an area near the position (each session 10 min). **e)** Air puff activation during acquisition (day 1) was similar between sham-control and aCB1KO mice (6.59 ± 0.73 vs. 6.13 ± 0.62, n = 17/15, p = 0.62, Mann-Whitney test). Comparing the activations binned into five-minute time intervals (small insets) revealed a significant decrease over time (n = 17/15, p < 0.001, two-way repeated measures ANOVA) without a difference between groups (n = 17/15, p = 0.64, two-way repeated measures ANOVA). On test day 2, no difference in the total transition number was found (1.41 ± 0.21 vs. 1.40 ± 0.34, n = 17/15, p = 0.73, Mann-Whitney test). After removal of the air puff from position A on day 3, the number of transitions at position A increased over time for both groups (small inset, sham control: 2.06 ± 0.66 vs. 3.94 ± 1.01, aCB1KO: 2.33 ± 0.70 vs. 4.93 ± 0.91, n = 17/15, time p = 0.00031, treatment p = 0.55, time x treatment p = 0.52, two-way repeated measures ANOVA) and the number of overall transitions was not significantly different between groups (6.00 ± 1.50 vs. 7.27 ± 1.45, n = 17/15, p = 0.45, Mann-Whitney test). **f)** Left panel: on day 4 (reversal), the air puff was moved to position B. Animals passed position A without a difference between groups (transitions, 3.47 ± 1.04 vs. 3.27 ± 1.02, n = 17/15, p = 0.77, Mann-Whitney test). aCB1KO mice activated the air puff at position B more frequently than sham control mice (3.65 ± 0.50 vs. 5.93 ± 0.61, n = 17/15, p = 0.0066, Student’s t-test). Right panel: comparing the number of air puff activations binned in five minutes intervals revealed a significant influence of time and experimental group (n = 17/15, time: p < 0.0001, treatment: p = 0.0066, time x treatment: p = 0.072, two-way repeated measure ANOVA). aCB1KO mice activated the air puff more often during the first 5 minutes compared to sham-inject mice (2.35 ± 0.33 vs. 4.07 ± 0.35, p = 0.0065, post-hoc Tukey test) but not during the last 5 minutes (1.29 ± 0.33 vs. 1.87 ± 0.35, p = 0.64, post-hoc Tukey test). Sham control mice did not show a difference over time (2.35 ± 0.33 vs. 1.29 ± 0.33, p = 0.13, post-hoc Tukey test) while aCB1KO mice did (4.07 ± 0.35 vs. 1.87 ± 0.35, p = 0.00063, post-hoc Tukey test). **g)** The number of animals activating the air puff at position B ≤ 3 times was significantly lower in aCB1KO mice (76.9 vs. 23.1 %, p = 0.036, Fisher’s exact test).

We next examined how sham injected and aCB1KO animals encode changes in the location of an aversive stimulus. We used a place avoidance test, in which mice first learn to avoid the location of an automatically triggered air puff at a specific position (**Fig. 6d**, position A) in an O-shaped maze for two days (day 1 and 2, probe trial on day 3). Subsequently, the aversive stimulus is switched to the opposite side of the O-maze (**Fig. 6d**, position B), requiring mice to encode the new aversive stimulus location. The spatial component in such tests is believed to be processed by the hippocampus whereas the integration of the aversive component requires the amygdala (Girardeau et al. 2017; Phillips and LeDoux 1992). Initially, both groups of animals rapidly learned to avoid the air puff (day 1 and 2) and reacted similarly to the removal of the air puff from position A on day 3 (**Fig. 6e-f**, **Suppl. Fig. 8a**). However, when the air puff was moved to position B on day 4, aCB1KO mice more often triggered the air puff in the beginning of the experiments (**Fig. 6f**) and avoided position B less than their control littermates (**Fig. 6g**, **Suppl. Fig. 8b-d**). Together these results indicate that aCB1KO animals do learn to avoid an aversive stimulus as efficiently as controls but display a transient deficit (**Fig. 6f**, right panel) when the spatial context needs to be updated.

## Discussion

A main hypothesis of our study was that dendritic spiking of CA1 pyramidal cells is controlled by the extracellular level of NMDAR co-agonists. Indeed, we found that exogenous D-serine and other manipulations of co-agonist levels transiently affected dendritic spiking in CA1 stratum radiatum so that an increase of extracellular co-agonist levels reduced the dendritic spike threshold and increased their amplitude, and vice versa. This indicates that the occupancy of the NMDAR co-agonist binding site dynamically determines NMDAR function and thereby dendritic integration and spiking.

Importantly, our study reveals that the activity of pyramidal cell populations controls dendritic spiking in a frequency-dependent manner via NMDAR co-agonist supply and astrocytic CB1Rs (see **Suppl. Fig. 9** for a schematic). Therefore, astrocytes close a positive feedback loop between pyramidal cell activity and their own dendritic spiking, which displays a bell-shaped frequency dependence with a peak at ~ 10 Hz. While pyramidal cell activity at 4 Hz did not affect dendritic spiking, stimulation at 10 Hz decreased the dendritic spike threshold and increased the dendritic spike amplitude. A straightforward explanation is that the dendrites of CA1 pyramidal cells are increasingly depolarized as the stimulation frequency increases and more efficiently activate NMDAR co-agonist release via CB1R-dependent astrocytic Ca^2+^ signaling, because the latter was shown to increase as the duration of direct neuronal depolarization is prolonged (Navarrete and Araque 2008). This raises the question why the effect on dendritic spiking decreases again at a stimulation frequency of 20 Hz and disappears at 40 Hz. Having excluded that the somatic firing of pyramidal cells cannot follow a 40 Hz stimulation, another possibility is that the frequency dependence is of dendritic origin. To test that hypothesis, we inhibited HCN-mediated currents, which are particularly strong in dendrites of CA1 pyramidal cells and confer a frequency-dependence to their excitability (Magee 1998; Narayanan and Johnston 2007). This prevented the modulation of dendritic spiking by pyramidal cell activity at 10 Hz and suggests that indeed dendritic excitability controls the modulation of dendritic spiking, because HCN expression by other neuronal subtypes is not relevant for our experimental approach. For instance, inhibition of HCN could hyperpolarize the dendritic trees of the stimulated pyramidal cell population thereby interfering with endocannabinoid release or inhibit dendritic resonance (Narayanan and Johnston 2007). In addition, the declining modulation of dendritic integration at higher frequencies could reflect a reduced propagation of somatic action potentials into the dendritic trees of stimulated cells because of increased dendritic inactivation of voltage-dependent sodium channels (Remy et al. 2009). Although the mechanisms underlying the frequency dependence clearly requires further dissection, our results demonstrate that extracellular NMDAR co-agonist levels are controlled by pyramidal cell activity via astrocytic CB1Rs with a bell-shaped frequency dependence peaking around the higher end of the theta frequency range.

Such a feedback loop could be relevant for the formation of spatial memories for several reasons. First, the investigated dendritic spiking has been implicated in the encoding of spatial information (see Introduction). Second, NMDARs participate in the generation of dendritic plateau potentials, which are driven by synaptic input from the entorhinal cortex and through CA3-CA1 synapses and contribute to place field formation (Bittner et al. 2015, 2017). Third, the observed peak of the dendritic spike modulation is in the theta range, which is an activity pattern that occurs for instance during spatial exploration (Vanderwolf 1969). Indeed, animals without astrocytic CB1Rs showed an impaired performance in an object location memory test. This reveals that astrocytic CB1Rs and NMDAR co-agonist signaling are important for encoding and/or storing object locations, in addition to their role in object recognition memory (Robin et al. 2018). Interestingly, the deliberate activation of astrocytes has recently been shown to play a role in contextual and object recognition memory too: Chemogenetic and optogenetic activation of hippocampal astrocytes improves the acquisition of contextual memory (Adamsky et al. 2018) and optogenetic activation of anterior cortical astrocytes improves remote object recognition memory (Iwai et al. 2021).

Further studying the role of astrocytic CB1Rs in spatial memory in a passive place avoidance test, we revealed that the spatial memory deficit uncovered in the object location test can be overcome if the relevance of a location is increased by an aversive stimulus, i.e., an air puff. A similar improvement of memory by aversive cues has been demonstrated previously. For instance, the performance of mice in the Morris water maze was increased in the presence of a predator odor, which was shown to involve the amygdala (Galliot et al. 2010). It is therefore likely that the air puff used in our spatial passive avoidance test enforces spatial learning by involving the amygdala. Interestingly, the spatial memory impairment of mice with CB1R-deficient astrocytes resurfaced transiently when the location of the aversive stimulus was moved (see **Fig. 6f**). In such experiments that test the association of an aversive stimulus and a context, the hippocampus is believed to encode the context (Phillips and LeDoux 1992). The transient deficit in acquiring the new location of the air puff is therefore likely of hippocampal origin and therefore associated with the impaired dendritic spiking of CA1 pyramidal cells in these mice in vitro. Whether this is in fact a causal relationship is a challenging experimental question. Another prediction arises from the fact that astrocytic CB1Rs mediate astrocytic Ca^2+^ transients and that its disruption impairs spatial memory. This indicates that some aspect of the spatial memory task such as the location of the animal should be encoded by astrocytic Ca^2+^ during exploration. Indeed, there is prominent astrocytic Ca^2+^ signaling during locomotion (Agarwal et al. 2017; Dombeck et al. 2007; King et al. 2020) and astrocytic activity has recently been used to predict an animal’s position in a familiar environment (Doron et al. 2021).

At level of NMDARs, our current results emphasize that the endogenous supply of the NMDAR co-agonist D-serine is dynamically regulated and that astrocytes and their Ca^2+^ signaling play an important role in setting the availability of D-serine (Adamsky et al. 2018; Henneberger et al. 2010; Papouin et al. 2017; Rasooli-Nejad et al. 2014; Robin et al. 2018; Takata et al. 2011). We demonstrate that astrocytic Ca^2+^ signals initiated by the activation of astrocytic CB1Rs (Navarrete and Araque 2008; Rasooli-Nejad et al. 2014; Robin et al. 2018) either pharmacologically or by endogenous pyramidal cell activity led to increased co-agonist levels and thereby a reduced dendritic spike threshold and an increased dendritic spike amplitude. This chain of events could be disrupted in experiments with a D-serine degrading enzyme, indicating that indeed D-serine had been released, and using CB1R-deficient astrocytes, which revealed that astrocytic CB1Rs are involved. Although we did not dissect the release mechanisms of D-serine, our results clearly demonstrate that astrocytes are key regulators of extracellular D-serine levels (Adamsky et al. 2018; Henneberger et al. 2010; Kronschläger et al. 2016; Panatier et al. 2006; Papouin et al. 2017; Rasooli-Nejad et al. 2014; Takata et al. 2011; Yang et al. 2003). In addition, we uncovered a previously unknown mechanism that dynamically adjusts NMDAR co-agonists levels according to neuronal activity.

These observations do not rule out that the NMDAR co-agonist glycine may also play a role for dendritic integration. A previous study has revealed that the co-agonist for synaptic NMDARs is primarily D-serine whereas it is glycine for extrasynaptic NMDARs (Papouin et al. 2012). The latter are however likely to be activated during glutamatergic synaptic crosstalk in the hippocampus (Scimemi et al. 2004), they contribute to NMDA spikes in cortical layer V pyramidal cell dendrites (Chalifoux and Carter 2011) and it is conceivable that activation of hippocampal extrasynaptic NMDARs becomes more prominent during the simultaneous activity of spatially clustered synapses that is required for triggering a local dendritic spike. Interestingly, extracellular glycine levels are actively maintained by glycine transporters in the hippocampus (Bergeron et al. 1998; Tsai et al. 2004) and we could previously demonstrate that they are modulated by neuronal activity using a newly-designed optical glycine sensor (Zhang et al. 2018). An interesting open question is therefore if activity-dependent changes of extracellular glycine levels also control dendritic spiking. Similarly, the spine-to-spine variability of glutamate uptake efficiency (Herde et al. 2020) and the increased glutamatergic crosstalk due to withdrawal of perisynaptic astrocyte processes from synapses undergoing long-term potentiation (Henneberger et al. 2020) may profoundly affect dendritic spike generation, their dependence on NMDAR co-agonists and place field formation.

## Acknowledgments

We thank Dr. Kurtulus Golcuk and Rebekka Zölzer for assistance with behavioral tests. Research was supported by the NRW-Rückkehrerprogramm (C.H.) and the German Research Foundation (DFG; SFB1089 B03, SPP1757 HE6949/1, FOR2795 HE6949/4, and HE6949/3 to C.H., SFB 1089 C04 and SPP 2041 to HB). We also acknowledge support by the Bonn Technology Campus.

## Author contributions

K.B. and C.H. designed, performed, and analyzed experiments involving iontophoretic glutamate application. N.M. and H.B. planned and carried out tests using glutamate uncaging. Ca^2+^ imaging was done by K.B. and E.M.S.. K.B. and K.H. performed additional electrophysiological tests. Behavioral tests were designed, performed and analyzed by K.B., A.N.H., A.Z., H.B., and T.O.. C.H. conceived the study, planned experiments, analyzed data, and wrote the manuscript, which was subsequently contributed to by all the authors.

## Methods

### Animals

All animals were housed under 12 h light/dark conditions and were allowed *ad libitum* access to food and water. The experiments were performed using 3-5-week-old Wistar rats (Charles River), 6-10-week-old male C57BL/6N wild type (Charles River) and transgenic mice of the following lines: GLAST-creERT2 (Mori et al. 2006), CB1^fl/fl^ (Marsicano et al., 2003), flox-top tdTomato (Madisen et al. 2010) (JAX 007908), flox-stop GCaMP5g-IRES-tdTomato (Gee et al. 2014) (JAX 024477). Transgenic mice of both genders were used to minimize breeding. To induce cre expression, 3-week-old mice were injected with tamoxifen (100 mg / kg BW; 1/day i.p., 5 days).

### Stereotactic injections

For stereotactic injections of rAAVs into the ventral CA1 region of the hippocampus, flox-stop GCaMP5g-IRES-tdTomato mice were deeply anesthetized (Fentanyl, Rotexmedica, 0.05 mg/kg bodyweight; Midazolam, Rotexmedica, 5.0 mg/kg bodyweight; Medetomidin, Cepitor from CPPharma, 0.5 mg/kg bodyweight; injection volume 0.1 ml/10 g bodyweight, i.p.). The eyes were covered with a crème (Bepanthen® eye and nose crème), and the anesthesia was confirmed by testing toe reflexes. Next, the head fur was shaved, the head was fixed in a stereotactic frame (Model 901, David Kopf Instruments) and the skin was locally anaesthetized (1 x 10 mg puff, Xylocain, Astra Zeneca). Then, a small incision was made, and the skull was carefully cleared from the remaining periosteum. A small whole (coordinates relative to *bregma*: anterior −3.5 mm, lateral −/+3 mm, ventral −2.5 mm) was drilled with a dental drill, and a beveled needle nanosyringe (nanofil 34G BVLD, WPI) was slowly inserted into the brain. After allowing the tissue to adjust, 0.5 μl of viral particles (2.94 x 10^13^ VP/ml, AAV1.CaMKII0.4Cre.SV40, V4567MI-R, PennCore) were injected under the control of a microinjection pump (100 nl/min, Micro4 Microsyringe Pump Controller, WPI). The needle was left in place for at least five minutes to avoid reflux into the needle track. Then, the needle was retracted slowly, and the procedure was repeated for the other hemisphere. Finally, the incision was sutured (anti-bacterial absorbable thread, Ethicon) and an antibiotic was applied (Refobacin 1mg/g, Gentamicin). The anesthesia was terminated (Naloxon, Puren, 1.2 mg/kg bodyweight; Flumazenil, Anexate from Hikma, 0.5 mg/kg bodyweight, Atipamezol, Antisedan from Ventoquinol, 2.5 mg/kg bodyweight; injection volume 0.1 ml/10 g bodyweight, i.p.) and the mouse was placed back into its home cage. Analgesia was applied 30 minutes before terminating the anesthesia as well as 8, 16 and 24 hours after the surgery (Buprenorphin, Buprenovet from Bayer, 0.05 mg/kg bodyweight; injection volume 0.1 ml/20g bodyweight, i.p.). These animals were used for experiments three to five weeks after surgery.

### Hippocampal slice preparation

Acute brain slices were prepared as previously described (Minge et al. 2017). Briefly, 300 μm slices were prepared in an ice-cold slicing solution (in mM: sucrose 105, NaCl 60, KCl 2.5, MgCl_2_ 7, NaH_2_PO_4_ 1.25, ascorbic acid 1.3, sodium pyruvate 3, NaHCO_3_ 26, CaCl_2_ 0.5 and glucose 10; osmolarity 300–310 mOsm/l). After a recovery of 15 minutes in 34°C-warm slicing solution, the slices were stored in artificial cerebrospinal fluid (ACSF, composition in mM: NaCl 131, KCl 2.5, MgSO_4_ 1.3, NaH_2_PO_4_ 1.25, NaHCO_3_ 21, CaCl_2_ 2 and glucose 10; pH 7.35–7.45; osmolarity 297–303 mOsm/l) for at least one hour before the start of the experiment. All experiments were conducted at ∼34°C.

### Glutamate iontophoresis

Acute slices were transferred into a submersion-type recording chamber mounted on a Scientifica two-photon excitation fluorescence microscope with a 40x/0.8 NA objective or 60x/1 NA objective (Olympus), placed onto a self-build grid and superfused with ACSF at 34° C. CA1 pyramidal cells were patched in the whole-cell configuration using a Multiclamp 700B amplifier (Molecular Devices) with borosilicate glass pipettes (3-5 MΩ resistance, GB150F-10, Science Products) filled with an intracellular solution containing in mM: KCH_3_O_3_S 135, HEPES 10, di-Tris-Phosphocreatine 10, MgCl_2_ 4, Na_2_-ATP 4, Na-GTP 0.4, Alexa Fluor 594 hydrazide 0.04 (Thermo Fisher Scientific), Fluo-4 pentapotassium salt 0.2 (Thermo Fisher Scientific), pH adjusted to 7.2 using KOH, osmolarity 290-295 mOsm/l. Cells were kept in the current clamp mode. Using a Ti:sapphire pulsed laser (Vision S, Coherent) the fluorescent indicators were excited (λ = 800 nm). A glutamate microiontophoresis system (MVCS-C-01C-150, NPI) was used for local dendritic activation. For this purpose, a borosilicate glass pipette (60-90 MΩ resistance, GB150F-10, Science Products), filled with 150 mM glutamic acid (pH adjusted to 7.0 with NaOH) and 50 μM Alexa Fluor 594 hydrazide (Thermo Fisher Scientific), was placed in close proximity (< 1 μm) to a spine on an apical oblique dendrite under the guidance of two-photon excitation fluorescence microscopy. We chose mostly dendrites directly branching off the apical trunk dendrite, which were stimulated at about 2/3 of their length from the apical trunk dendrite. Leakage of glutamate was prevented by a small, positive retain current (< 8 nA). The iontophoretic stimulation duration was adjusted (< 0.8 ms) so that reliable somatic excitatory postsynaptic potentials (EPSPs), dendritic spikes and action potentials (APs) could be recorded with increasing stimulation intensity (25 to 50 pA steps until the occurrence of an AP, 3 s interstimulus interval). All experiments were conducted in the presence of 50 μM picrotoxin (abcam Biochemicals) to inhibit GABA_A_ receptors. Only cells with an initial access resistance (R_a_) < 20 MΩ and with < 20 % change during the time course of the recording were included in the analysis.

### Glutamate uncaging

Acute slices were transferred to a submerged recording chamber on a dual galvanometer based scanning system (Prairie Technologies). CA1 pyramidal cells were visualized with infrared oblique illumination optics and a water immersion objective (60x, 0.9 NA, Olympus) and somatic whole-cell current clamp recordings were performed with a BVC-700 amplifier (Dagan Corporation). Data were filtered at 10 kHz and sampled at 50 kHz with a Digidata 1440 interface controlled by pClamp Software (Molecular Devices). Patch-pipettes were pulled from borosilicate glass (outer diameter 1.5 mm, inner diameter 0.8 mm; Science Products) with a P-97 Puller (Sutter Instruments) to resistances of 2 to 5 MΩ in bath and whole-cell series resistances ranging from 8 to 30 MΩ. The standard internal solution contained (in mM): 140 K-gluconate, 7 KCl, 5 HEPES, 0.5 MgCl2, 5 phosphocreatine, 0.16 EGTA. Internal solutions were titrated to pH 7.3 with KOH, had an osmolality of 295 mOsm/l, and contained 100 μM Alexa Fluor 594 (Thermo Fisher Scientific). Voltages were not corrected for the calculated liquid-junction potential of +14.5 mV. The membrane potential was adjusted to −75 mV for all recordings. Cells with unstable input resistances or lacking overshooting action potentials were discarded as well as recordings with holding currents >-200 pA at −60 mV and access resistances > 30 MOhm. For two-photon glutamate uncaging at apical oblique dendrites of CA1, MNI-caged-L-glutamate 15 mM (Tocris) was dissolved in HEPES-buffered solution (in mM as follows: 140 NaCl, 3 KCl, 1.3 MgCl_2_, 2.6 CaCl_2_, 20 D-glucose, and 10 HEPES, pH 7.4 adjusted with NaOH, 305 mOsm/l) and applied using positive pressure via glass pipettes (< 1 MΩ) placed in close proximity to the selected apical oblique dendrites of CA1 neurons. We used two ultrafast laser beams of Ti:sapphire pulsed lasers (Chameleon Ultra, Coherent), which were tuned to 860 nm to excite the Alexa 594 and to 720 nm to photo-release glutamate at 10-15 dendritic spines of a dendritic segment of ~ 10 μm in length. The intensity of each laser beam was independently controlled with electro-optical modulators (Conoptics Model 302RM). MNI-glutamate was uncaged at an increasing number of spines (2-15) with 0.5 ms exposure times and the uncaging spot was rapidly moved from spine to spine with a transit time of ~0.1 ms. The laser power at the slice surface was kept below 22 mW to avoid photo damage. Glutamate was uncaged onto a sequence of single spines to evoke excitatory post synaptic potentials (uncaging evoked EPSPs, uEPSPs). To quantify deviations from linearity in dendritic integration, the arithmetic sum of individual uEPSPs of single spines (expected EPSP) was compared to the measured EPSP evoked by rapid sequential glutamate uncaging onto the same set of spines. The rate of rise of the dendritic spike’s initial fast phase was calculated from the maximum dV/dt value. The amplitude of the slow phase of dendritic spikes was obtained from the same dendritic spikes used to quantify dV/dt. The dendritic spike threshold was calculated as the amplitude of the expected EPSP at which dendritic spikes first occurred. All data analyses were done with Clampfit 9.2, (Molecular Devices) and IGOR Pro (Wavemetrics).

### Alveus stimulation

Selective activation of axons in the alveus was achieved through an incision between CA1 and the subiculum that spared the alveus. Next to the subicular cut, a clustered bipolar stimulation electrode (CE2F75, FHC) was placed (**Suppl. Fig. 2a**). A borosilicate glass pipette (2-4 MΩ resistance, GB150F-10, Science Products) filled with ACSF was positioned at the boarder of stratum oriens and stratum pyramidale of CA1 to record the evoked population spikes. The stimulation intensity was adjusted to 80% of that evoking the maximum population spike amplitude. Only slices with a minimum population spike amplitude of 0.8 mV were used for experiments. In a subset of experiments the glass pipette was removed and a CA1 pyramidal cell within the same area (± 70 μm) was patched and stimulated using glutamate iontophoresis as described above. In another set of recordings, extracellular single unit responses were recorded. In these recordings, a borosilicate glass pipette (5-7 MΩ resistance, GB150F-10, Science Products) was placed into the pyramidal cell layer during 10 Hz alveus stimulation to blindly identify single unit responses. These responses are characterized by a sharp upward reflection followed by a smaller downward reflection. Next, recordings at different frequencies (4, 10 and 40 Hz) of alveus stimulation were done in quadruplets. A maximum of three single unit recordings per slice were performed.

### Field recordings in acute hippocampal slices

In a subset of experiments (**Suppl. Fig. 6a-b**), field EPSPs (fEPSPs) were evoked by electrical stimulation of CA3-CA1 Schaffer collaterals (100 μs) and recorded in the CA1 stratum radiatum through an extracellular pipette (patch pipette as described above) filled with the extracellular solution. For the characterization of synaptic transmission, the stimulus intensity was increased in a stepwise manner from 20 to 640 μA. For paired-pulse experiments, the stimulus intensity was set to obtain a half-maximal response and the interstimulus interval was varied. In another subset of experiments (**Suppl. Fig. 6c**), an extracellular recording electrode was placed in the pyramidal cell layer to record the population spike after synaptic stimulation. Again, the stimulus intensity was increased in a stepwise manner from 20 to 640 μA to compare population spikes and their dependence on the strength of stimulation between experimental groups.

### Pharmacology

The following drugs in the respective concentrations were used: 5 μM AM251 (abcam), 0.17 U/ml DAAO (Sigma), 50 μM D-APV (abcam), 10 μM D-serine (Sigma), 10 μM NBQX (abcam), 0.5 μM Ro 25-6981 (abcam), 1 μM WIN55,212-2 mesylate (Sigma), 10 μM ZD7288 (Tocris).

### Astrocyte whole-cell patch clamp and Ca^2+^ imaging

Experiments were performed as described previously (Anders et al. 2014; Henneberger and Rusakov 2012). Acute slices were transferred to a submersion-type recording chamber mounted on a Scientifica two-photon excitation fluorescence microscope with a 40x/0.8 NA objective (Olympus). A stratum radiatum astrocyte at a depth of 40 - 60 μm was recorded from in the whole-cell patch clamp configuration using a Multiclamp 700B amplifier (Molecular Devices) using borosilicate glass pipettes (3.0-3.5 MΩ resistance, GB150F-10, Science Products) using an intracellular solution containing in mM: KCH_3_O_3_S 135, HEPES 10, di-Tris-Phosphocreatine 10, MgCl_2_ 4, Na_2_-ATP 4, Na-GTP 0.4, Alexa Fluor 594 hydrazide 0.04 (Thermo Fisher Scientific), Fluo-4 pentapotassium salt 0.2-0.4 (Thermo Fisher Scientific), pH adjusted to 7.2 using KOH, osmolarity 290-295 mOsm/l and kept in the current-clamp mode. Fluorescent dyes were excited with a Ti:sapphire pulsed laser (Vision S, Coherent, λ = 800 nm). The cellular identity was confirmed by their membrane potential (< −80 mV), their low input resistance (< 5 MΩ), their typical morphology and gap junction coupling. Recordings in which the access resistance exceeded 30 MΩ were excluded from the analysis.

In experiments from **Fig. 2**, time-lapse frame scanning (0.75-1.5 Hz) was performed after sufficient dye diffusion (> 20 min) during baseline and after bath application of 10 μM of WIN 55,212-2. A subset of these experiments was performed in the presence of the inverse endocannabinoid receptor agonist AM251 (5 μM). For analysis, the fluorescence intensity ratio (R, Fluo-4 / Alexa 594, both channels corrected for background fluorescence) was quantified in a region of interest sparing the soma and major branches during WIN55 application (R) and before (R_0_). The effect of WIN55 was quantified by calculating R/R_0_.

In experiments from **Fig. 3**, alveus stimulation was combined with astrocyte whole-cell patch clamp recordings and Ca^2+^ imaging. An alveus stimulation electrode was positioned, the stimulation intensity was set as described and an astrocyte was patched. After sufficient dye diffusion (> 20 min), time lapse imaging was performed during baseline, during alveus stimulation (3x 20 Hz for 1 s), and after bath application of the inverse endocannabinoid receptor agonist AM251 (5 μM) during a second round of alveus stimulation. The overall change of astrocytic Ca^2+^ signals was estimated by first calculating the ratio of both fluorescent indicators (R, Fluo-4/Alexa Fluor 594, botch channels corrected for background fluorescence) in one or six regions of interest covering the astrocytic territory sparing the soma and major branches (Fiji, Image J). For each step of the recording (baseline, alveus stimulation, alveus stimulation in AM251), the resting value R_0_ and the average ΔR (change relative to R_0_) were determined and ΔR/R_0_ was calculated as a measure of overall Ca^2+^ signal activity. ΔR/R_0_ during stimulation/treatment was compared to the baseline period to quantify the effect of stimulation/treatment (**Fig. 3c**, right panel). ROIs that showed an increase of ΔR/R_0_ of more than 5% during alveus stimulation were defined to be responders (**Fig. 3c**, left panel) and used to analyze the effect of AM251.

### Calcium imaging using genetically encoded Ca^2+^ indicators

The expression of GCaMP5g and tdTomato in astrocytes or CA1 pyramidal cells was established through crossbreeding flox-stop GCaMP5g-IRES-tdTomato mice with GLAST-creERT2 mice and intraperitoneal tamoxifen injections (astrocytes) as described before (King et al. 2020) or rAAV injections (pyramidal cells, see above). Horizontal hippocampal slices were prepared and recorded from on an Olympus FV10MP microscope with a 25x/1.05NA objective or a Scientifica two-photon excitation fluorescence microscope with a 40x/0.8 NA objective at 34 °C in ACSF (supplemented with 50 μM Picrotoxin as in all other acute slice experiments). Using a Ti:sapphire pulse laser (Vision S, Coherent) both fluorescent proteins were excited (λ = 910 nm). Acquisition of the images was performed using ScanImage Software or Olympus Fluoview 4.2.

For experiments in **Supp. Fig. 2** on CA1 pyramidal cells, the response of up to six CA1 pyramidal cells to 20 Hz alveus stimulation was determined by line scanning (~2 ms per line, 1500 lines). The intensity profile of the fluorescent ratio (GCaMP5g/tdTomato, background corrected) was analyzed using Fiji (Image J). Recordings were performed in quadruplets for at least 16 cells per slice to determine the percentage of responding cells. An arbitrary threshold of 25% well above the noise level was defined to identify responding cells.

In experiments from **Fig. 4** and **Suppl. Fig. 4**, astrocytic calcium transients were recorded using time-lapse frame scanning (128 x 128 pixels, 80 μm x 80 μm, 2.96 Hz) during baseline and 10 Hz or 40 Hz alveus stimulation for three minutes each. The overall changes of astrocytic Ca^2+^ signals were estimated by first calculating the ratio of both fluorescent indicators (R, GCaMP5g/tdTomato, both channels corrected for background fluorescence) in regions of interest covering the astrocytic territory sparing the soma and major branches (Fiji, Image J). For each period of the recording (baseline, alveus stimulation at 10 or 40 Hz), the resting value R_0_ and the average ΔR (change relative to R_0_) were determined and ΔR/R_0_ was calculated as a measure of overall Ca^2+^ signal activity. ΔR/R_0_ during stimulation was compared to the baseline period to quantify the effect of stimulation (**Fig. 4c**). ROIs that showed an increase of ΔR/R_0_ of more than 5% during alveus stimulation were defined to be responders (**Fig. 4b**). The same data set was also analyzed by manual identification of Ca^2+^ transients and manual definition of regions of interest (3 μm x 3 μm) using Fiji (Image J) (**Suppl. Fig. 4**). The frequency of events during the baseline period and alveus stimulation was calculated (**Suppl. Fig. 4b-c**). For the analysis of Ca^2+^ signal amplitudes, the fluorescence ratio of both Ca^2+^ indicators (R, GCaMP5g/tdTomato, both channels corrected for background fluorescence) was determined immediately before each event (R_0_) and its change (ΔR) at the peak to calculate its amplitude relative to R_0_ (ΔR/R_0_). Mean amplitudes of astrocytic Ca^2+^ transients were then compared between baseline recordings and alveus stimulation (**Suppl. Fig. 4d**).

### Immunohistochemistry

Mice were anesthetized using an overdose of ketamine and xylazine, the chest cavity opened and transcardially perfused with at least 20 ml ice-cold paraformaldehyde (4% in PBS). After removal of the brain, it was placed in the same solution overnight for post-fixation. Next, the brain was cut in 50 μm thick, horizontal slices using a vibratome (Leica). After blocking for 2h in 10 % normal goat serum (NGS, Merck Millipore) and 0.5 % Triton (AppliChem) in PBS, the slices were stained with primary antibodies for GFAP (polyclonal rabbit anti-cow GFAP, 1:500; Z0334, Dako) and NeuN (monoclonal mouse anti NeuN, 1:200; MAB377, Chemicon) in PBS with 5 % NGS and 0.1 % Triton overnight at 4°C. After washing, the slices were places in in PBS with 2 % NGS, 0.1 % Triton and the secondary antibodies for anti-rabbit (polyclonal goat anti-rabbit IgG (H+L)-Alexa Fluor 488, 1:500; A11034, Ivitrogen) and anti-mouse (polyclonal goat anti-mouse IgG (H+L)-biotin, 1:500; 115-065-003, Dianoca) for one to one and a half hours at room temperature. This was followed by a streptavidin conjugate to fluorescently tag biotin (Alexa Fluor 647 streptavidin conjugate, 1:600; S32357, Thermo Fisher Scientific). For nuclear staining, the slices were placed in water with 0.5 % Hoechst (33342, ThermoFisher Scientific) for ten minutes at room temperature. Finally, the slices were mounted onto microscope slides (Thermo Scientifica) with a ProLong Gold antifade mountant (Invitrogen) and stored at 4°C. Image stacks of the hippocampal CA1 area (512 x 512 μm, 450 x 450 pixel, pixel size 1.14 μm, 2 μm z-steps) were taken using a laser scanning confocal microscope (TCS SP8, Leica). Three sections of 20 μm depth each were averaged per animal for analysis. Cells were counted by the nuclear staining using Fiji (Image J).

### Behavioral tests

GLAST-creERT2 mice (Mori et al. 2006) were crossed with CB1^fl/fl^ mice (Marsicano et al. 2003) and flox-stop tdTomato mice (Madisen et al. 2010). After weening, mice of both genders were injected with either tamoxifen or a sham solution and housed under a reversed 12h light/dark conditions. Handling and behavioral experiments as well as tamoxifen injection and analysis were performed by individual, blinded experimenters. Behavioral testing was performed in a separate, quiet area during the beginning of the dark phase. The room was lit with dimmed red light and spatial cues were present around the arena. All test and training sessions were recorded using a Basler acA1300-200um camera system mounted above the arena (1280 x 1024 pixels). After each individual animal test, the arena was carefully cleaned using 70 % ethanol.

Habituation (5 days) and the open field test (first day of habituation) took place in a rectangle arena (40 x 60 cm) that was constructed of dark gray PVC with 23 cm wall height and light-gray flooring. The mice were tracked automatically by three points (head, body, tail), manually checked and if needed adapted using EthoVision XT14 (Noldus). For analysis of anxiety-related behavior an outer zone of an eight cm wide gallery and an inner zone of 24 x 44 cm were defined.

The same arena was used for testing object location memory. Two identical objects (4 cm in diameter) were placed randomly in the corners of the arena with a distance of 9 cm from the walls. The mice were placed in the middle of the arena and allowed to explore the objects freely for ten minutes. After a 24h delay, one of the objects was placed to a different corner and the mice were placed back into the arena for five minutes of exploration. Tracking and analysis was performed with EthoVision XT14 (Noldus). Object exploration was defined as the nose-point being within 2 cm of the object, with body orientation towards the object. The discrimination index ((time at object with novel location - time at object with constant location) / total time) was used as an indicator of the object location memory. Animals that showed excessive climbing or little exploration (exploration time <5 s in 10 min or <2 s in 5 min; total distance <1 m in 5 min) as well as outliers for the analyzed parameters (mean ± 2SD) were excluded (in total n = 5).

Spatial working memory was investigated using a Y-maze with three identical arms, each 40 cm long and 8 cm wide, interconnected at 120° angle. Each arm had a different spatial cue at the end. The mice were placed in the middle of the arena and were allowed to explore it freely for ten minutes. Tracking (one point, body) and analysis was performed with EthoVision XT14 (Noldus). The number and order of entrances into the second half of an arm was determined automatically. The number of performed alternations (triplet of visits to all three arms) was divided by the number of possible alternations to calculate the alternation index. Animals were considered to have an intrinsic arm preference or aversion when they entered an arm 10 % more or less often than expected (33%, relative arm visits < 23% or > 43%). Those animals were excluded from further analysis (n = 3).

The passive place avoidance test was performed in a modified open field arena with a 10 cm wide gallery. Light sensors on each short side of the gallery (position A and B) were used to activate an air puff (one bar, nitrogen gas) when the animal was disrupting the light bridge. The animal was placed in the long arm of the arena in equal distance to the possible air puff locations. Each day a single trial of free exploration (10 minutes) was performed. The animal’s behavior was tracked using EthoVision XT14 (Noldus). Activation of the light bridge was determined through entry of the nose-point into a small area (1 cm wide) at the air puff location.

### Statistics

Statistical analyses were performed with OriginPro (OriginLab) and Matlab (Mathworks). Data are expressed and displayed as mean ± s.e.m. (standard error of mean) or in box plots. In the latter, the box indicates the 25th and 75th, the whiskers the 5th and 95th percentiles, the horizontal line in the box the median and the mean is represented by a filled circle. Statistical tests were performed after investigating whether data followed a normal distribution (Shapiro-Wilk test). Depending on the outcome, statistical tests were performed using non-parametric (Kruskal-Wallis test, Friedman test, Mann-Whitney test, Wilcoxon signed rang test) or parametric (ANOVA, repeated measures ANOVA, Student’s t-tests) approaches as indicated. In graphs, statistical significance is indicated by asterisk. * for p < 0.05, ** for p < 0.01 and *** for p < 0.001. Please see Suppl. Table 2 for a comprehensive overview of statistical tests and their results.

