## Supplementary material for "An astrocytic signaling loop for frequency-dependent control of dendritic integration and spatial learning"

### Supplementary figure 1

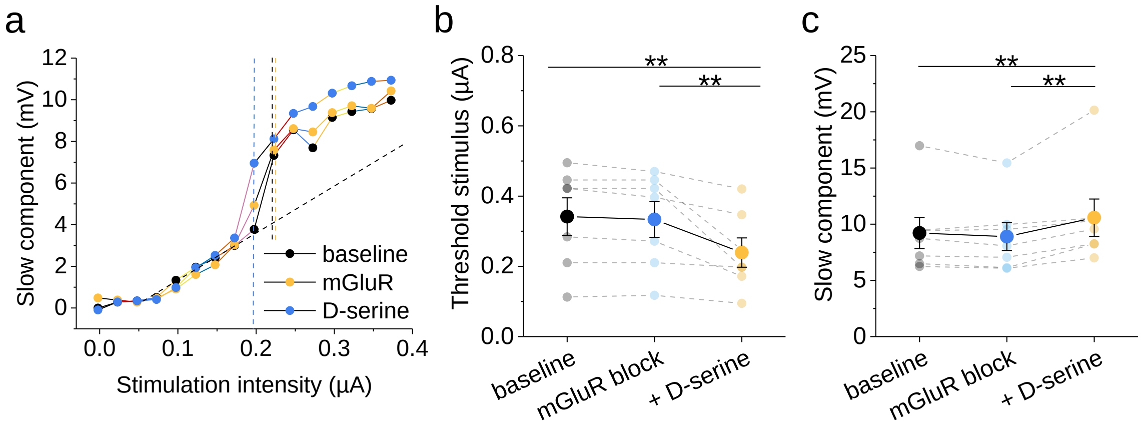


**Supplementary figure 1: Activity of metabotropic glutamate receptors (mGluRs) does not control dendritic spikes evoked by iontophoretic glutamate application.**

**a)** Sample experiment probing dendritic spiking under baseline conditions (black), in the presence of the mGluR inhibitors LY341495 (50 µM) and MPEP (10 µM), which at these concentrations should block all relevant mGluRs, and after additional wash-in of D-serine (10 µM).

**b)** Threshold stimulus of evoking a dendritic spike (0.34 ± 0.05 µA vs. 0.33 ± 0.05 µA vs. 0.24 ± 0.04 µA from left to right, n = 7, $F(1.03,6.19) = 9.50$, $p = 0.020$, one-way repeated-measures ANOVA; post-hoc Fisher LSD: baseline vs. mGluR $t(12) = 0.31$, $p = 0.76$, baseline vs. D-serine $t(12) = 3.92$, $p = 0.0020$, mGluR vs. D-serine $t(12) = 3.61$, $p = 0.0036$)

**c)** Amplitude of slow component of the dendritic spike (9.22 ± 1.39 mV vs. 8.89 ± 1.24 mV vs. 10.58 ± 1.66 mV, n = 7, $F(1.08,6.47) = 10.59$, $p = 0.015$ one-way repeated measures ANOVA; post-hoc Fisher LSD: baseline vs. mGluR $t(12) = 0.83$, $p = 0.42$, baseline vs. D-serine $t(12) = 3.51$, $p = 0.0043$, mGluR vs. D-serine $t(12) = 0.433$, $p = 0.00097$)

**Supplementary figure 2**

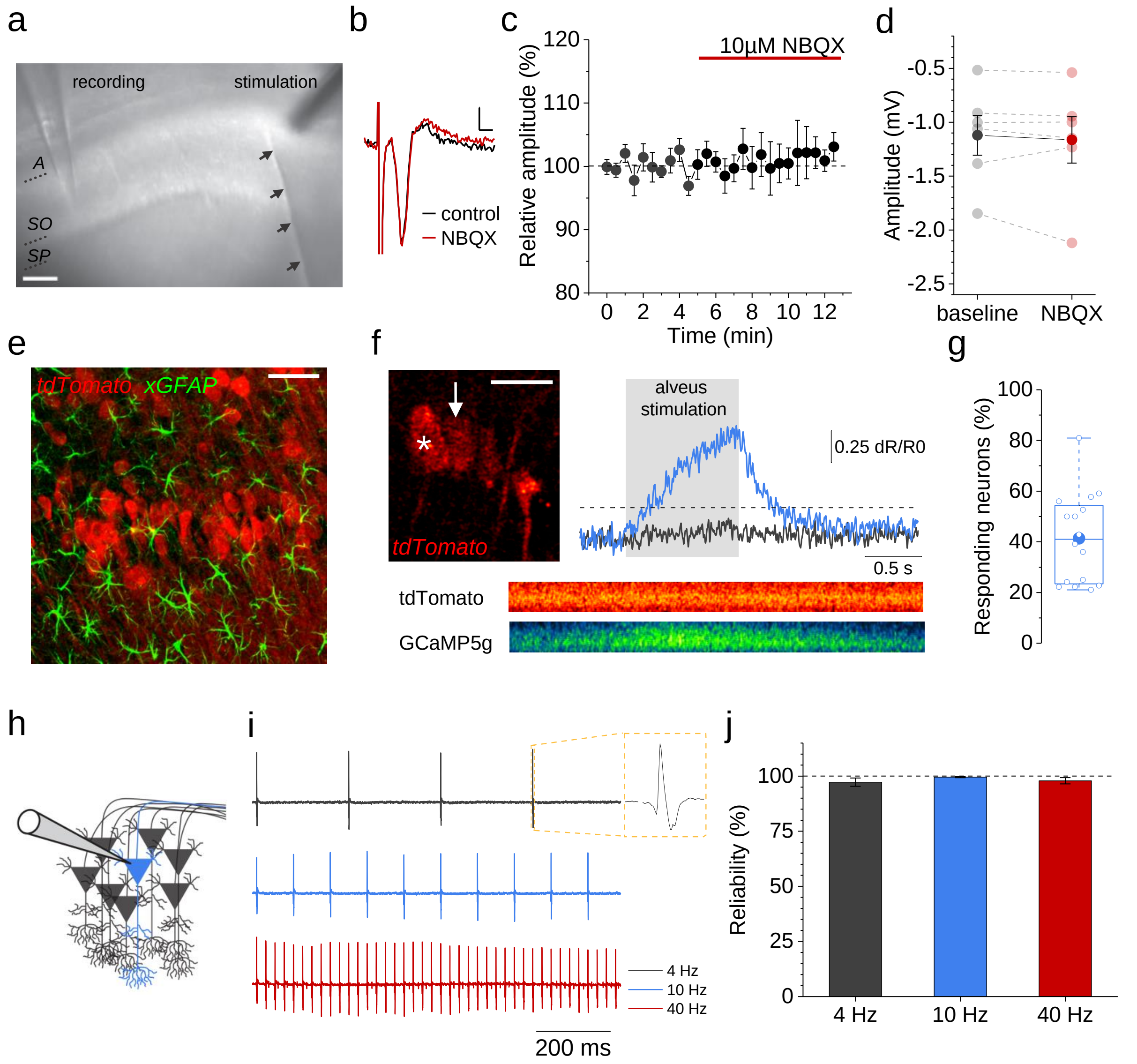


**Supplementary figure 2: Experimental approach for retrogradely evoking CA1 pyramidal cell activity by alveus stimulation.**

**a)** Placement of extracellular recording and stimulation electrodes in the CA1 pyramidal cell layer (SP) and alveus (A), respectively (SO, stratum oriens). Arrows indicate position of a cut that was made to isolate retrograde activation of CA1 pyramidal cells by stimulating their axons.

**b)** Example of a population spike evoked by stimulation in the alveus (scale bar, 0.1 mV, 1 ms)

**c-d)** The amplitude of the evoked population spike is stable over time (**c**) and independent of excitatory synaptic transmission (d, population spike amplitude, control -1.12 ± 0.18 mV, NBQX -1.16 ± 0.21 mV, n = 6, t(5) = 0.76, p = 0.48, paired Student's t-test)

**e)** Virally induced recombination in CA1 pyramidal cells using a CaMKII-cre AAV in a mouse line with cre-dependent expression of GCaMP5g and tdTomato (Gee et al. 2014). Staining for GFAP and tdTomato revealed no recombination in astrocytes (confocal microscopy).

**f)** CA1 pyramidal cells expressing tdTomato in an acute slice (top left panel, scale bar 20 µm). Line scans through several somata were performed during stimulation of the alveus (examples in bottom panels, width of line scan 8.5 µm, length 2129 ms). Top right panel: representative filtered traces of the ratio of GCaMP5g and tdTomato (background-subtracted and normalized to baseline) are illustrated for a nonresponding (asterisk, grey trace) and a responding neuron (arrow, blue trace). Cells were considered responders if the ratio increased more than 25 % over baseline (dashed line).

**g)** On average, 41.3 ± 4.4 % of the cells in an acute brain slice (n = 16) responded to alveus stimulation.

**h-i)** Cartoon illustrating extracellular single unit recordings and examples for different frequencies of alveus stimulation.

**j)** The reliability of evoking single unit activity did not depend on alveus stimulation frequency. When a pyramidal cell responded to single stimuli, several trains of stimuli were delivered at three frequencies (4, 10 and 40 Hz) and the probability of observing a unit spike was calculated. No significant differences were detected (4 Hz: 97.3 ± 1.9 %, 10 Hz: 99.5 ± 0.3 %, 40 Hz: 97.9 ± 1.4, n = 16 cells from 7 independent experiments, $F(1.01,15.13) = 1.55$, $p = 0.23$ one-way repeated-measures ANOVA).

**Supplementary figure 3**

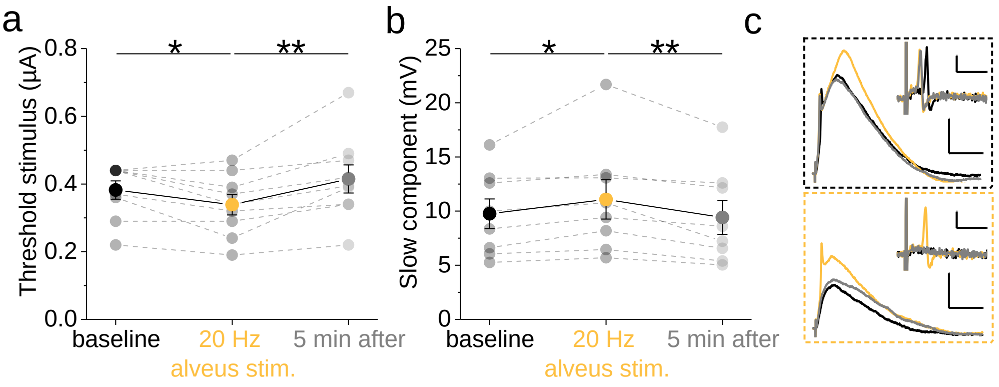


**Supplementary figure 3: Changes of dendritic spikes induced by alveus stimulation are reversible.** The threshold stimulus of dendritic spikes and their slow component were analyzed in baseline conditions, during 20 Hz alveus stimulus and 5 minutes after alveus stimulation.

**a)** Threshold stimulus during baseline, 20 Hz alveus stimulation, and five minutes later (baseline: 0.38 ± 0.03 µA, 20 Hz: 0.34 ± 0.03 µA, 5 min: 0.41 ± 0.04 µA, n = 9, $\chi^2(2) = 9.06$, p = 0.011 Friedman test; post-hoc Wilcoxon signed-rank tests: z = 2.12, p = 0.031 for baseline vs. 20 Hz, z = 1.26, p = 0.21 for baseline vs. 5 min, z = 2.61, p = 0.0039 for 20 Hz vs. 5 min)

**b)** Analysis of the slow component of dendritic spike of the same recordings. n = 8 because somatic spikes prevented analysis of the slow component in one recording (baseline: 9.75 ± 1.37 mV, 20 Hz: 11.08 ± 1.81 mV, 5 min: 9.41 ± 1.56, n = 8, F(2,14) = 5.80, p = 0.015 one-way repeated measures ANOVA; post-hoc Fisher's LSD test, t(14) = 2.57, p = 0.022 for baseline vs. 20 Hz, t(14) = 0.66, p = 0.52 for baseline vs. 5 min, t(14) = 3.22, p = 0.0061 for 20 Hz vs. 5 min).

**c)** Sample traces for baseline (black), 20 Hz alveus stimulation (orange) and 5 min after (grey). The upper panel (black border) shows traces for all three conditions recorded with the stimulus intensity that just evoked a dendritic spike under baseline conditions (threshold stimulus). Note that during alveus stimulation the slow component is increased. The lower panel (orange border) shows traces for all three conditions recorded with the stimulus intensity that just evoked a dendritic spike during 20 Hz alveus stimulation. Because alveus stimulation reduces the threshold stimulus, no dendritic spikes occur at that simulation intensity under baseline conditions and 5 minutes after alveus stimulation. Insets display dV/dt with the same color coding. Scale bars 2 mV and 20 ms and in insets 2 mV/ms and 5 ms.

### Supplementary figure 4

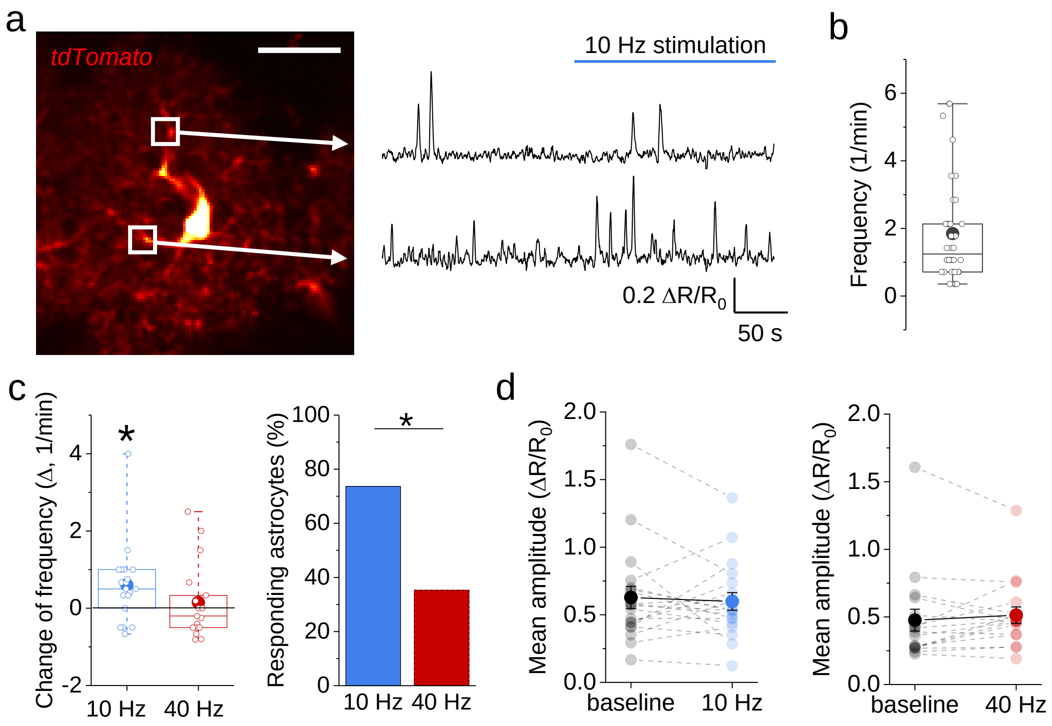


**Supplementary figure 4: Frequency-dependent activation of astrocytes by alveus stimulation.** $Ca^{2+}$ transients were monitored in astrocytes expressing GCaMP5g and tdTomato. Additional analyses of the data presented in Fig. 4a-c.

**a)** Example of an astrocyte expressing tdTomato (left panel, scale bar 20 µm) and GCaMP5g (not shown). Right panel: examples of astrocytic $Ca^{2+}$ transients from the two regions of interest (ROI) illustrated in the left panel. The fluorescence ratio (R) of GCaMP5g and tdTomato (R) was calculated and its changes (ΔR) were normalized to its resting value ($R_0$). See Methods for further details.

**b)** Frequencies of $Ca^{2+}$ transients during baseline recordings before alveus stimulation in active (1.85 ± 0.25 events per minute, n = 36 astrocytes).

**c)** Change of $Ca^{2+}$ transient frequency during alveus stimulation at 10 and 40 Hz (blue and red, respectively). Alveus stimulation at 10 but not 40 Hz significantly increased the frequency of $Ca^{2+}$ transients (left panel; 10 Hz: 0.60 ± 0.23, n = 19, z = 2.45, p = 0.011; 40 Hz: 0.15 ± 0.24, n = 17, z = 0.17, p = 0.86; one-population Wilcoxon signed-rank tests). The percentage of astrocytes showing an increase in frequency was significantly higher with 10 Hz stimulation than with 40 Hz (right panel; p = 0.043, Fisher's exact test).

**d)** Alveus stimulation did not affect the amplitude of astrocytic $Ca^{2+}$ transients (10 Hz, left panel: 0.63 ± 0.08 vs. 0.60 ± 0.07, n = 19, z = 0.46, p = 0.65; 40 Hz, right panel: 0.48 ± 0.08 vs. 0.51 ± 0.06, n = 17, z = 1.09, p = 0.28; Wilcoxon signed rank tests).

**Supplementary figure 5**

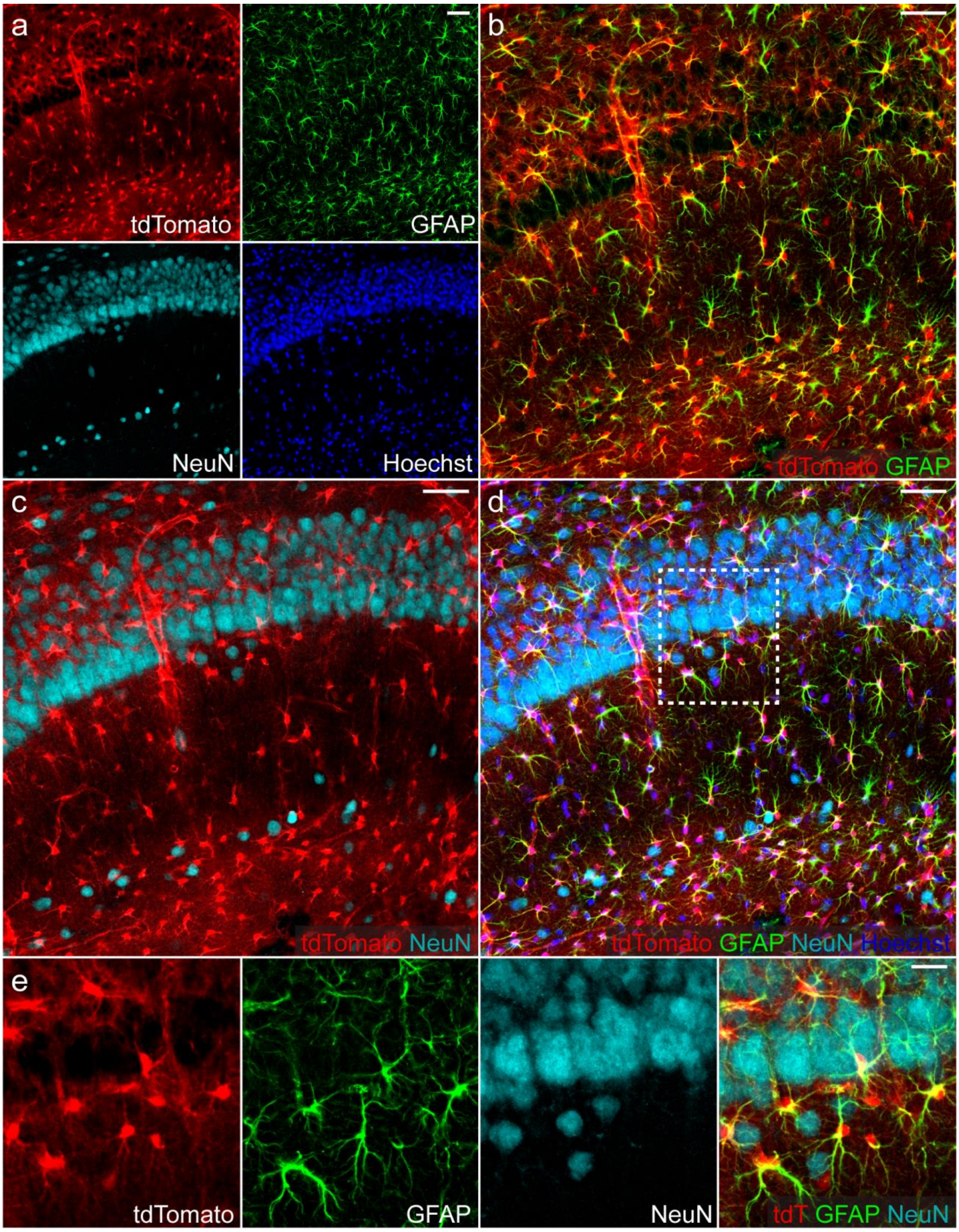


**Supplementary figure 5: Characterisation of astrocyte-specific recombination after tamoxifen-injection in GLASTcreERT2 mice in the CA1 region**. GLASTcreERT2 (Mori et al. 2006) mice crossed with a flox-stop tdTomato reporter line (Madisen et al. 2010) were injected with tamoxifen and their hippocampal CA1 region characterized by immunohistochemistry and confocal microscopy. Scale bars are 50 µm in **a-d** and 20 µm in **e**.

**a)** Examples of tdTomato expression (top left panel), astrocytic GFAP expression (top right panel), neuronal NeuN expression (bottom left panel) and DNA label Hoechst to visualize cell nuclei (bottom right panel)

**b)** The vast majority of tdTomato-expressing cells is positive for the astrocytic marker GFAP (overlay of tdTomato and GFAP). Of all tdTomato-expressing cell, 89.7 ± 1.4 % (n = 3) are positive for GFAP. Of all GFAP-positive cells, 98.7 ± 0.8 % (n = 3) express tdTomato.

**c)** Neuronal NeuN expression and tdTomato expression do not overlap (overlay of tdTomato and NeuN). Of all NeuN-positive cells, only 2.0 ± 1.8% (n = 3) were also positive for tdTomato.

**d)** Overlay of tdTomato, GFAP, NeuN and Hoechst. A higher magnification of the delineated area is shown in **e**.

**Supplementary figure 6**

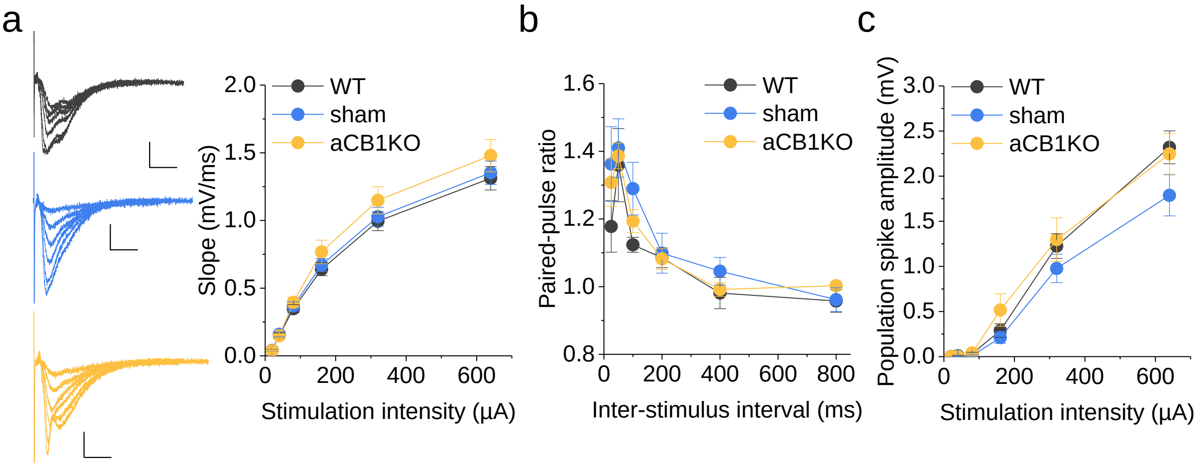


**Supplementary figure 6: Basic properties of CA3-CA1 synaptic transmission are not altered by the absence of astrocytic CB1 receptors.** Comparison of fEPSPs in CA1 stratum radiatum evoked by electrical stimulation of CA3-CA1 synapses in slices from wild-type animals (WT)´and GLASTcreERT2 x CBR1 $^{fl/fl}$ mice injected with tamoxifen (aCB1KO) or a control solution (sham).

**a)** Stimulus-response curve. Examples are shown on the left (top: WT, black; middle: sham, blue; bottom: aCB1KO, yellow; scale bars: 10 ms and 0.5 mV). Right panel: summary of experiments (WT: n = 49 from 19 animals; sham: n = 41 from 16 animals; aCB1KO: n = 46 from 17 animals; two-way repeated measures ANOVA, between groups, $F(2,133) = 0.80$, $p = 0.37$).

**b)** The paired-pulse ratio of fEPSP was recorded at several inter-stimulus intervals. No statistically significant differences were found between groups (WT: n = 7 from 4 animals, sham: n = 14 from 5 animals; aCB1KO: n = 14 from 6 animals; two-way repeated measures ANOVA, between groups, $F(2,30) = 0.77$, $p = 0.39$)

**c)** Population spikes were recorded from the CA1 pyramidal layer and evoked by stimulation of CA3-CA1 synaptic connections at various stimulations intensities. No statistically significant differences between groups (WT: n = 46 from 20 animals, sham: n=32 from 17 animals; aCB1KO: n = 30 from 14 animals; two-way repeated measures ANOVA, between groups, $F(2,105) = 1.59$, $p = 0.21$).

### Supplementary figure 7

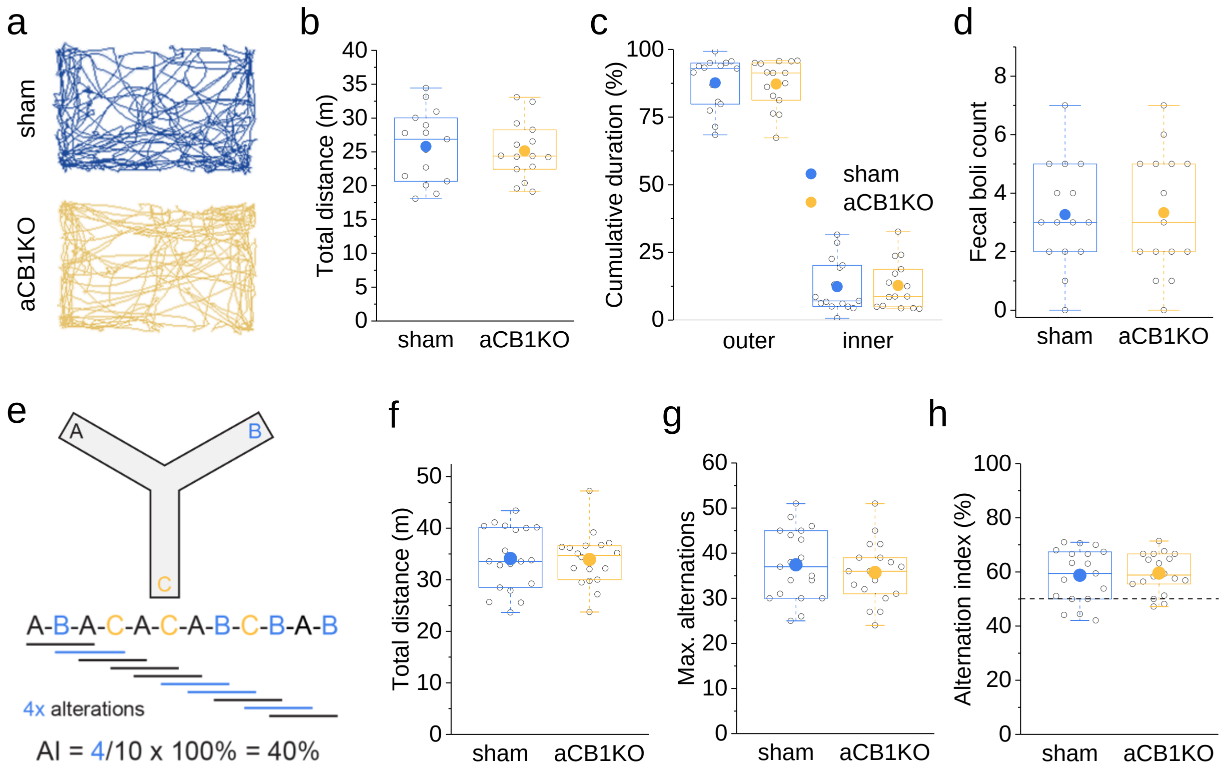


**Supplementary figure 7: Absence of astrocytic CB1Rs does not affect the behaviour of mice in an open arena or in a spontaneous alteration test (Y-maze).** Comparison of GLASTcreERT2 x CBR1 $^{fl/fl}$ mice injected with tamoxifen (aCB1KO) or a control solution (sham).

**a)** Examples of tracks of mice in an open arena (top: sham, blue; bottom: aCB1KO, yellow).

**b-d)** The two groups of animals (n = 15 both groups) did not differ significantly regarding the total distance travelled in the arena (**b**, sham: 25.8 ± 1.4 m, aCB1KO: 25.1 ± 1.1 m, t(28) = 0.37, p = 0.71, Student's t-test), the cumulative time spent in the outer part (less than 8 cm from arena border) or inner part of the arena (**c**, outer, sham: 87. 7 ± 2.5 %, outer, aCB1KO: 87.2 ± 2.3 %, inner, sham: 12.3 ± 2.5 %, inner, aCB1KO 12.8 ± 2.3 %, two-way repeated measurement ANOVA: for groups F(1,28) = 1.00, p = 0.33, for interaction F(1,28) = 0.02, p = 0.90, for inner/outer F(1,28) = 492.49, p < 0.0001) or the number of fecal boli count as an indirect measure of anxiety (d, fecal boli count, sham: 3.3 ± 0.5, aCB1KO: 3.3 ± 0.5, t(28) = 0.09, p = 0.93, Student's t-test).

**e)** Schematic representation of the spontaneous alteration test (Y-maze) with arms A, B and C. The alternation index (AI) is calculated as the percentage of fully alternating triple entries (blue lines) out of all sequential triplets (blue and black lines).

**f-g)** The two groups of animals (sham: n = 19, aCB1KO: n = 18) differ not significantly regarding the total distance travelled (**f**, sham: 34.1 ± 1.4 m, aCB1KO: 33.9 ± 1.2 m, t(35) = 0.10, p = 0.92, Student's t-test), the maximum number of alternations (**g**, sham: 37.4 ± 1.8, aCB1KO: 35.7 ± 1.6, t(35) = 0.71, p = 0.48, Student's t-test) or the alternation index (sham: 58.8 ± 2.2, aCB1KO: 59.6 ± 1.7 %, t(35) = 0.29, p = 0.78, Student's t-test).

**Supplementary figure 8**

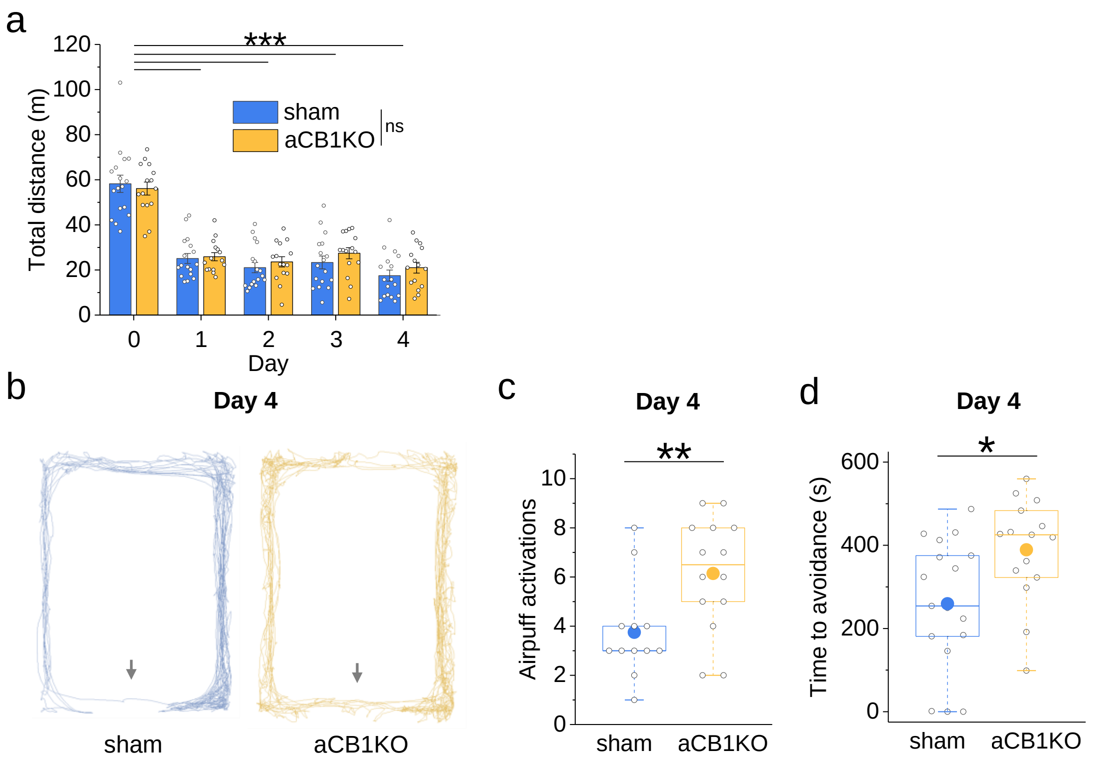


**Supplementary figure 8: Additional analyses of place avoidance learning in control mice (sham) and mice without astrocytic CB1 receptors (aCB1KO).** See **Fig. 6** for experiment and main findings. For animal model also see **Suppl. Fig. 4-5**.

**a)** The total distance travelled by mice declined rapidly during the experiment but did not differ between experimental groups (two-way repeated measures ANOVA; for days $F(4,120) = 186.20$, $p < 0.001$, post-hoc Tukey test $p < 0.001$ as indicated by asterisks; for groups $F(1,30) = 0.32$, $p = 0.58$; for interaction $F(4,120) = 1.23$, $p = 0.30$; n = 17 and 15 for sham and aCB1KO, respectively).

**b)** Sample tracks at day 4. The new air puff location is marked by arrows. Note that the sham-treated animals successfully and rapidly avoid the air puff.

**c)** Number of air puff activations in both groups on day 4. In contrast to Fig. 6, animals that did not fully explore the arena on day 3 (no air puff in any location) were excluded from this analysis, because they had no opportunity to notice the absence of an air puff on day 3. This did not change the results qualitatively. Sham-treated animals activated fewer air puffs, i.e., avoided the new air puff area more successfully (sham: 3.75 ± 0.57, n = 12; aCB1KO: 6.14 ± 0.62, n = 14; $t(24) = 2.82$, $p = 0.0096$, Student's t-test).

**d)** How quickly animals avoided the new air puff location was quantified by measuring the time between the first triggered air puff and the last, after which the airpuff area was completely avoided. This time to avoidance is low when mice rapidly acquire the new location of the aversive stimulus. Sham-treated animals displayed a significantly shorter time to avoidance (sham: 259.6 ± 38.2, n = 17; aCB1KO: 389.2 ± 32.3, n = 15; $t(30) = 2.55$, $p = 0.016$, Student's t-test).

**Supplementary figure 9**

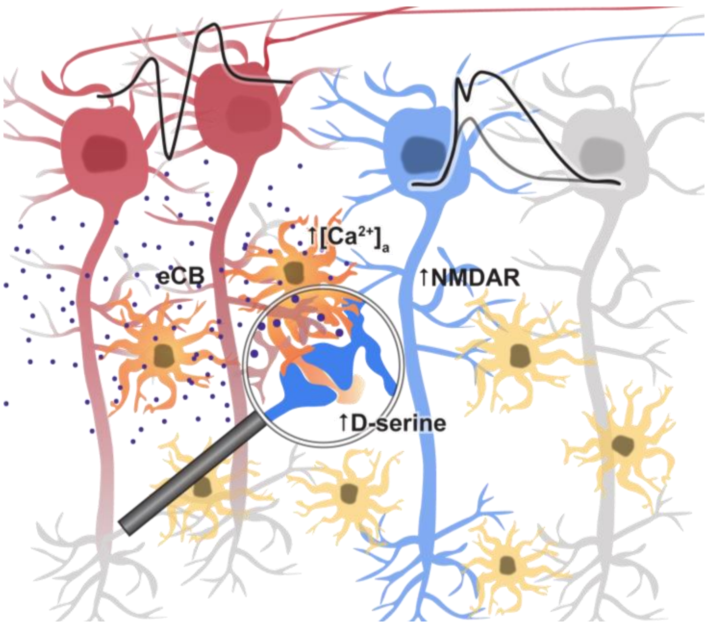


**Schematic of astrocytic mechanism promoting supralinear dendritic integration:** Active pyramidal cells (red) release endocannabinoids, which act on astrocytic CB1 receptors and thereby increase extracellular D-serine levels. This leads to increased opening of NMDA receptors and a lower threshold of dendritic spikes and a higher NMDA component. This positive feedback loop is primarily engaged by theta-like activity of pyramidal cells. Its disruption leads to deficits of spatial memory.

**Supplementary table 1**

| parameter | wildtype (mean ± SEM) | sham (mean ± SEM) | aCB1KO (mean ± SEM) | n | p | F value |
|---|---|---|---|---|---|---|
| **Rm (MΩ)** | 136.8 ± 14.1 | 119.3 ± 9.7 | 132.8 ± 10.5 | 7/8/9 | 0.54 | F(2,23) = 0.63 |
| **Vm (mV)** | -63.4 ± 2.96 | -66.9 ± 1.95 | -67.7 ± 1.18 | 7/8/9 | 0.21 | F(2,23) = 1.70 |
| **AP threshold (mV)** | -43.1 ± 2.11 | -50.0 ± 1.88 | -48.9 ± 1.09 | 9/10/10 | 0.02 | F(2,28) = 4.42 |
| **AP amplitude (mV)** | 93.1 ± 1.7 | 93.7 ± 4.0 | 94.1 ± 1.4 | 8/10/10 | 0.97 | F(2,27) = 0.03 |
| **AP upstroke (mV/ms)** | 409.3 ± 16.8 | 407.2 ± 9.2 | 400.1 ± 14.7 | 9/9/10 | 0.87 | F(2,27) = 0.12 |

**Supplementary table 1. Properties of CA1 pyramidal cells in wildtype, sham-injected mice and mice with deletion of astrocytic CB1 receptors (aCB1KO).** Also see Supplementary figures 4 and 5). The following parameters were analyzed in whole-cell patch clamp and field recordings: the membrane resistance ($R_M$), the resting membrane potential ($V_M$), the action potential (AP) threshold, AP amplitude, AP maximum depolarization rate (upstroke) and the population spike amplitude (Pop. spike) in alveus stimulation experiments and comparted using one-way ANOVA. Post-hoc analysis for AP threshold (Tukey): q = 3.95, p = 0.025 for wildtype vs. sham, q = 3.31, p = 0.068 for wildtype vs. aCB1KO, q = 0.65, p = 0.89 for sham vs. aCB1KO.

### Supplementary table 2

| Figure | parameter | | n | \|test\| | p value |
|---|---|---|---|---|---|
| 1d | threshold stimulus | | 16 | t(15) = 5.37 | 0.000077 |
| | slow component | | 16 | t(15) = 8.91 | 0.00000022 |
| 1e | threshold stimulus | | 16 | t(15) = 5.21 | 0.00011 |
| | slow component | | 14 | t(13) = 5.00 | 0.00024 |
| 1i | threshold | | 6 | t(5) = 2.78 | 0.039 |
| | slow component | | 6 | t(5) = 4.07 | 0.0097 |
| 1j | threshold | | 6 | t(5) = 2.59 | 0.049 |
| | slow component | | 6 | t(5) = 2.65 | 0.045 |
| 2b | $R/R_0$ | control | 13 | t(12) = 3.38 | 0.0054 |
| | | in AM251 | 5 | t(4) = 0.71 | 0.52 |
| 2c | threshold stimulus | | 9 | t(8) = 3.24 | 0.012 |
| | slow component | | 8 | t(7) = 2.62 | 0.034 |
| 2d | threshold stimulus | | 6 | t(5) = 0.17 | 0.87 |
| | slow component | | 6 | t(5) = 0.50 | 0.64 |
| 3c | $R/R_0$ relative to baseline | | 9 | t(8) = 3.04 | 0.016 |
| 3d (S3a) | threshold stimulus | 4 Hz | 5 | t(4) = 0.00 | 0.99 |
| | | 10 Hz & 40 Hz | 8 | F(2,14) = 17.4 | 0.00016 |
| | | *post-hoc 10 Hz* | | t(14) = 5.56 | 0.000071 |
| | | *post-hoc 40 Hz* | | t(14) = 1.05 | 0.31 |
| | | *post-hoc 10 Hz vs. 40 Hz* | | t(14) = 4.50 | 0.00050 |
| | | 20 Hz & 5 min after (S3a) | 9 | $\chi^2(2) = 9.06$ | 0.011 |
| | | *post-hoc 20 Hz* | | z = 2.12 | 0.031 |
| | | *post-hoc 5 min* | | z = 1.26 | 0.21 |
| | | *post-hoc 20 Hz vs. 5 min* | | z = 2.61 | 0.0039 |
| | change | 4 Hz, 10 Hz, 20 Hz & 40 Hz | 5/8/9/8 | F(3,26) = 5.51 | 0.0046 |
| | | *post-hoc 4 Hz vs. 10 Hz* | | t = 3.67 | 0.0011 |
| | | *post-hoc 10 Hz vs. 20 Hz* | | t = 1.95 | 0.049 |
| | | *post-hoc 10 Hz vs. 40 Hz* | | t = 3.17 | 0.0039 |
| 3d (S3b) | slow component | 4 Hz | 5 | t(4) = 1.40 | 0.23 |
| | | 10 Hz & 40 Hz | 8 | F(2,14) = 19.30 | 0.000095 |
| | | *post-hoc 10 Hz* | | t(14) = 5.75 | 0.00005 |
| | | *post-hoc 40 Hz* | | t(14) = 0.84 | 0.41 |
| | | *post-hoc 10 Hz vs. 40 Hz* | | t(14) = 4.91 | 0.00069 |
| | | 20 Hz & 5 min after (S3b) | 8 | F(2,14) = 5.80 | 0.015 |
| | | *post-hoc 20 Hz* | | t(14) = 2.57 | 0.022 |
| | | *post-hoc 5 min* | | t(14) = 0.66 | 0.52 |
| | | *post-hoc 20 Hz vs. 5 min* | | t(14) = 3.22 | 0.0061 |
| | change | 4 Hz, 10 Hz, 20 Hz & 40 Hz | 5/8/8/8 | $\chi^2(3) = 8.77$ | 0.032 |
| | | *post-hoc 4 Hz vs. 10 Hz* | | z = 1.98 | 0.045 |
| | | *post-hoc 10 Hz vs. 40 Hz* | | z = 2.36 | 0.015 |
| 3f | threshold stimulus | | 7 | t(6) = 0.66 | 0.53 |
| | slow component | | 7 | t(6) = 0.20 | 0.85 |
| 3g | threshold stimulus | | 8 | t(7) = 0.35 | 0.73 |
| | slow component | | 8 | t(7) = 1.20 | 0.27 |

| Figure | parameter | | | n | \|test\| | p value |
|---|---|---|---|---|---|---|
| **4c** | $\Delta R/R_0$ relative to baseline | | | 26/25 | z = 4.31 | 0.0000046 |
| **4d** | threshold stimulus | 10 Hz in ZD7288 | | 8 | t(7) = 1.05 | 0.33 |
| **4d** | threshold stimulus | WIN55 in ZD7288 | | 10 | t(9) = 2.68 | 0.025 |
| **4d** | change | 10 Hz control, 10 Hz in ZD7288 & WIN55 in ZD7288 | | 8/8/10 | $\chi^2(2) = 14.06$ | 0.00088 |
| **4d** | change | *post-hoc 10 Hz control & 10 Hz in ZD7288* | | 8/8/10 | z = 3.10 | 0.00062 |
| **4d** | change | *post-hoc 10 Hz control & WIN55 in ZD7288* | | 8/8/10 | z = 1.02 | 0.32 |
| **4d** | change | *post-hoc 10 Hz & WIN55 in ZD7288* | | 8/8/10 | z = 3.15 | 0.00055 |
| **4e** | slow component | 10 Hz in ZD7288 | | 8 | t(7) = 0.67 | 0.52 |
| **4e** | slow component | WIN55 in ZD7288 | | 10 | t(9) = 4.77 | 0.0010 |
| **4e** | change | 10 Hz control, 10 Hz in ZD7288 & WIN55 in ZD7288 | | 8/8/10 | $\chi^2(2) = 10.53$ | 0.0052 |
| **4e** | change | *post-hoc 10 Hz control & 10 Hz in ZD7288* | | 8/8/10 | z = 2.68 | 0.0047 |
| **4e** | change | *post-hoc 10 Hz control & WIN55 in ZD7288* | | 8/8/10 | z = 0.49 | 0.63 |
| **4e** | change | *post-hoc 10 Hz & WIN55 in ZD7288* | | 8/8/10 | z = 2.80 | 0.0031 |
| **5c** | threshold stimulus | WT | | 7 | t(6) = 5.36 | 0.0017 |
| **5c** | threshold stimulus | sham | | 8 | t(7) = 5.74 | 0.00071 |
| **5c** | threshold stimulus | aCB1KO | | 9 | z = 0 | 1 |
| **5c** | change | WT, sham & aCB1KO | | 7/8/9 | F(2,21) = 11.74 | 0.00038 |
| **5c** | change | *post-hoc WT & sham* | | 7/8/9 | 0.41 | 0.68 |
| **5c** | change | *post-hoc WT & aCB1KO* | | 7/8/9 | 4.26 | 0.00035 |
| **5c** | change | *post-hoc sham & aCB1KO* | | 7/8/9 | 3.98 | 0.00067 |
| **5d** | slow component | WT | | 7 | t(6) = 3.11 | 0.021 |
| **5d** | slow component | sham | | 8 | t(7) = 3.95 | 0.0056 |
| **5d** | slow component | aCB1KO | | 9 | t(8) = 1.66 | 0.14 |
| **5d** | change | WT, sham & aCB1KO | | 7/8/9 | F(2,21) = 9.11 | 0.0014 |
| **5d** | change | *post-hoc WT & sham* | | 7/8/9 | t = 0.84 | 0.41 |
| **5d** | change | *post-hoc WT & aCB1KO* | | 7/8/9 | t = 3.04 | 0.0062 |
| **5d** | change | *post-hoc sham & aCB1KO* | | 7/8/9 | t = 4.05 | 0.00058 |
| **6b** | total exploration | | | 10/12 | z = 0.63 | 0.54 |
| **6c** | discrimination index | | | 10/12 | t(20) = 2.86 | 0.0098 |
| **6e** | transition position A | day 1 | | 17/15 | z = 0.50 | 0.62 |
| **6e** | transition position A | day 2 | | 17/15 | z = 0.36 | 0.73 |
| **6e** | transition position A | day 3 | | 17/15 | z = 0.76 | 0.45 |
| **6e** | transition position A | inset day 1 | time | 17/15 | F(1,30) = 139.71 | < 0.00001 |
| **6e** | transition position A | inset day 1 | treatment | 17/15 | F(1,30) = 0.22 | 0.64 |
| **6e** | transition position A | inset day 3 | time | 17/15 | F(1,30) = 16.60 | 0.00031 |
| **6e** | transition position A | inset day 3 | treatment | 17/15 | F(1,30) = 0.36 | 0.55 |
| **6e** | transition position A | inset day 3 | interaction | 17/15 | F(1,30) = 0.43 | 0.52 |
| **6f** | transitions | position A | | 17/15 | z = 0.29 | 0.77 |
| **6f** | transitions | position B | | 17/15 | t(30) = 2.92 | 0.0066 |
| **6f** | transitions | inset | time | 17/15 | F(1,30) = 28.42 | 0.0000092 |
| **6f** | transitions | inset | treatment | 17/15 | F(1,30) = 8.51 | 0.0066 |
| **6f** | transitions | inset | interaction | 17/15 | F(1,30) = 3.48 | 0.072 |
| **6f** | transitions | inset | *post-hoc 1st time bin* | 17/15 | t(30) = 5.04 | 0.0066 |
| **6f** | transitions | inset | *post-hoc 2nd time bin* | 17/15 | t(30) = 1.68 | 0.64 |
| **6f** | transitions | inset | *post-hoc sham* | 17/15 | t(30) = 3.21 | 0.13 |
| **6f** | transitions | inset | *post-hoc aCB1KO* | 17/15 | t(30) = 6.27 | 0.00063 |

**Supplementary table 2. Detailed information on statistics.**
